# A genomic portrait of zebrafish transposable elements and their spatiotemporal embryonic expression

**DOI:** 10.1101/2021.04.08.439009

**Authors:** Ni-Chen Chang, Quirze Rovira, Jonathan N. Wells, Cédric Feschotte, Juan M. Vaquerizas

## Abstract

There is considerable interest in understanding the effect of transposable elements (TEs) on embryonic development. Studies in humans and mice are limited by the difficulty of working with mammalian embryos, and by the relative scarcity of active TEs in these organisms. Zebrafish is an outstanding model for the study of vertebrate development and over half of its genome consists of diverse TEs. However, zebrafish TEs remain poorly characterized. Here we describe the demography and genomic distribution of zebrafish TEs and their expression throughout embryogenesis using bulk and single-cell RNA sequencing data. These results reveal a highly dynamic genomic ecosystem comprising nearly 2,000 distinct TE families, which vary in copy number by four orders of magnitude and span a wide range of ages. Longer retroelements tend to be retained in intergenic regions, whilst short interspersed nuclear elements (SINEs) and DNA transposons are more frequently found nearby or within genes. Locus-specific mapping of TE expression reveals extensive TE transcription during development. While two thirds of TE transcripts are likely driven by nearby gene promoters, we still observe stage and tissue-specific expression patterns in self-regulated TEs. Long terminal repeat (LTR) retroelements are most transcriptionally active immediately following zygotic genome activation, whereas DNA transposons are enriched amongst transcripts expressed in later stages of development. Single-cell analysis reveals several endogenous retroviruses expressed in specific somatic cell lineages. Overall, our study provides an important resource for using zebrafish as a model to study the impact of TEs on vertebrate development.

## Introduction

TEs are selfish genetic elements that replicate and mobilize within host genomes. They have colonized all vertebrate species sequenced to date but with differential success, accounting for between 4 and 60% of their genomes (Sotero-Caio et al., 2017). The success of TEs is dependent on their propagation through the germline. Thus, the time and place in which they are active is critical to their long-term survival in host genomes. Undifferentiated embryonic cells are one of the ‘niches’ adopted by TEs that facilitate their propagation (Haig, 2016). Whilst the mobility of TEs is thought to be generally deleterious to the host, the accumulation of TEs in the genome represents a source of raw genetic material that may be coopted during evolution to benefit diverse cellular functions, including functions related to embryogenesis (Durruthy-Durruthy et al., 2016; Garcia-Perez et al., 2016; Jachowicz et al., 2017; Lu et al., 2014; Wang et al., 2014). Zebrafish, a powerful model organism to study embryonic development, is also notable for its very high TE and repetitive DNA content (53%, Howe et al., 2013). As yet, however, little is known about the TE ecosystem of the zebrafish genome. Are TE families uniformly distributed across the genome or do they preferentially accumulate in certain regions? What is the demographic profile of zebrafish TEs? Does the diversity of zebrafish TE families result in distinct spatial and temporal patterns of expression during development? Are these expression patterns related to the intrinsic properties of individual TEs or are they driven by their genomic locale? In this work, we aim to answer these questions in order to establish the groundwork for the study of TEs in zebrafish development.

TEs exploit a variety of transcriptional and translational mechanisms to expand in the host genome. Based on their transposition intermediates, TEs are classified as retrotransposons or DNA transposons (Finnegan, 1989; Wells and Feschotte, 2020). Retrotransposons reverse-transcribe their own RNA and then insert the DNA copy back into the genome. Most retrotransposons carry internal promoters with cis-regulatory sequences that recruit the host transcriptional machinery to drive their own expression, much like host genes (Bowen, 2003; Brind’Amour et al., 2018; Faulkner et al., 2009; Robbez-Masson and Rowe, 2015; Romanish et al., 2007). In contrast, most DNA transposons directly excise themselves and re-insert elsewhere in the host genome, a process mediated by transposase genes encoded by autonomous DNA transposons (Fricker and Peters, 2014; Hickman and Dyda, 2016; Spradling et al., 2011). Compared to retrotransposons, the mechanisms directing the expression of DNA transposons are generally less characterized. Some do contain promoter sequences but these tend to be weak and not cell type-specific (Palazzo et al., 2017, 2019). Additionally, the diversity and relative abundance of DNA transposons and retrotransposons varies widely across species. For example, although TEs comprise nearly half of both human and zebrafish genomes, retrotransposons account for ~95% of all TEs in the human genome, but only ~10% in zebrafish (Howe et al., 2013; Lander et al., 2001). In contrast, approximately 40% of the zebrafish genome comprises of DNA transposons, while in humans they occupy just ~3% (Pace and Feschotte, 2007). Overall, all the major lineages of eukaryotic transposons, including the rarer types, can be found within the zebrafish genome, which harbors a much greater diversity of TEs than is typically observed in mammalian genomes (Chalopin et al., 2015; Furano et al., 2004; Howe et al., 2013).

Genome-wide studies have revealed that TEs are expressed in a tightly regulated fashion during mammalian embryonic development. In human and mouse early embryos, TE transcripts comprise up to 15% of the transcriptome (Göke et al., 2015; Peaston et al., 2004; Svoboda et al., 2004). While the expression pattern and regulatory activities of TEs during development likely reflect how they have exploited distinct cellular niches to propagate, these activities may also be integrated in normal developmental programs. For example, the expression of the murine long interspersed nuclear element-1 family (LINE1) can be detected shortly after fertilization and peaks at the 2-cell stage in mouse embryos, while cells are still totipotent (Fadloun et al., 2013; Peaston et al., 2004). This expression not only promotes LINE1 transposition in mouse early embryos (Richardson et al., 2017), but provocatively, it may also be essential for proper embryonic development (Jachowicz et al., 2017; Percharde et al., 2018). Endogenous retroviruses (ERVs), which are affiliated with LTR retrotransposons, are also transcriptionally active in a highly stage-specific manner in mammalian embryos (Göke et al., 2015; Grow et al., 2015). The expression of MERVL – a murine-specific ERV family – peaks at the 2-cell stage of embryogenesis and contributes to the expression of more than 50 chimeric MERVL-host gene transcripts in the mouse embryo (Macfarlan et al., 2012; Peaston et al., 2004). Similarly, HERVH, a primate-specific family, is specifically expressed from the 8-cell to the blastocyst stage and marks cells with higher pluripotent potential (Fort et al., 2014; Göke et al., 2015; Wang et al., 2014, 2016). Thus, understanding TE expression is important not only for understanding the biology of TEs, but also that of the host. However, most of what we know about the transcriptional activity of vertebrate TEs during embryogenesis comes from studies conducted in human or mouse, which harbor a very limited diversity of TEs relative to other vertebrates, and indeed most animals (Wells and Feschotte, 2020).

Little has been reported about the expression of TEs in zebrafish, but a few families have been serendipitously identified as markers of specific stages of embryonic development. For example, BHIKHARI, a zebrafish ERV family, is expressed exclusively in the mesendoderm lineage during gastrulation (Chen and Schier, 2001; Vogel and Gerster, 1999). A distantly related ERV, BHIKHARI-2 (also known as *crestin*) was discovered as a specific marker of the neural crest (Luo et al., 2001; Rubinstein et al., 2000). Despite these intriguing observations, BHIKHARI elements have not been characterized further and there is a general dearth of information regarding the genomic characteristics and expression of individual TE families in zebrafish. Previous studies examining zebrafish TEs on a genome-wide scale have been limited to broad patterns at the level of TE classes or subclasses (e.g. LTR, LINE, etc.) (Chalopin et al., 2015; Gao et al., 2016; Yang et al., 2020). However, different TE families within the same TE class can behave very differently when it comes to their genomic distribution or expression patterns (Feschotte et al., 2002; Ishiuchi et al., 2015; Rodriguez-Terrones and Torres-Padilla, 2018; Stitzer et al., 2019).

To establish a foundation for future work on the activity of TEs in zebrafish embryogenesis, we have performed a detailed characterization of the genomic landscape and embryonic expression of zebrafish TEs. Our study highlights the staggering diversity of TEs in zebrafish, yields insights into the effect of selection on the genomic distribution of different TE types and describes a wide diversity of transcriptional patterns through early development.

## Results

### The genomic landscape of zebrafish TEs

Using RepeatMasker to annotate the *Danio rerio* reference genome (GRCv11), we mapped the location of sequences related to a total of 1931 non-redundant TE families catalogued in Dfam and Repbase (Bao et al., 2015; Storer et al., 2021). These families include representatives of all major classes and subclasses of eukaryotic TEs, including LTR, non-LTR (LINEs and SINEs) and tyrosine recombinase retroelements, as well as DDE-type DNA transposons, rolling-circle elements (RC, i.e. Helitrons), Mavericks/Polintons and Cryptons (Wells and Feschotte, 2020; Wicker et al., 2007). Collectively, interspersed TEs account for 59.5% of the genome, with DNA transposons accounting for 46.2% and retroelements 13.2% (Supp. Data 1). Note that these values are higher than previously reported, likely as a result of improvements in the quality of the zebrafish reference genome since its initial publication (Howe, 2020; Howe et al., 2013). Amongst retroelements, the genome proportion of LTRs, LINEs and SINEs is 6.0%, 4.1% and 3.1% respectively, while tyrosine recombinase-mediated retroelements (DIRS and Ngaro superfamilies) account for 2.1%. DNA transposons are dominated by DDE-type transposons, which comprise 43.5% of the genome, whereas the more exotic Helitrons, Cryptons and Maverick/Polinton elements make up 1.3%, 0.9% and 0.5% respectively.

### DNA transposons tend to be older and more abundant than retroelements

We estimated the age of TEs by generating phylogenetic trees for all families with at least 10 copies of at least 100 base pairs in length (n=1880, Supp. Data 2), and then calculated the median length of terminal branches for each family (measured in nucleotide substitutions per site). This measure correlates well with estimates calculated using divergence from family consensus sequences, but avoids biases caused by family sub-structure (Supp. Fig. 1, Stitzer et al., 2019). Based on the presence of many families with identical insertions across the genome (i.e. branch length = 0) we can infer that all of the major TE classes in zebrafish - with the possible exception of SINEs, of which there are only 14 annotated families - contain either recently, or currently active families (Supp. Data 3).

Using this measure of age, we observed a moderate positive correlation between the average age of TE families and their copy number (Fig. 1, Spearman’s ρ=0.57, p≈0). There are very few examples of low copy-number elements that are also old; for example, of families with fewer than 50 copies, just three have a median branch length greater than 0.1 substitutions per site. In contrast, there are 45 very young families (fewer than 0.05 substitutions per site on average) with more than 1000 copies, and it is therefore likely that there are many families transpositionally active in zebrafish populations. We also observe significant differences in age and copy-number between TE classes: DNA transposon families are typically older and present at higher copy-number than both LINE and LTR retroelement families (Fig. 1). This trend could indicate either a recent increase in the rate of activity of retroelements relative to DNA transposons, or differences in the rate at which DNA transposons and retrotransposons are fixed in the population or deleted after insertion (Frahry et al., 2015; Kapusta et al., 2017).

**Figure 1.**
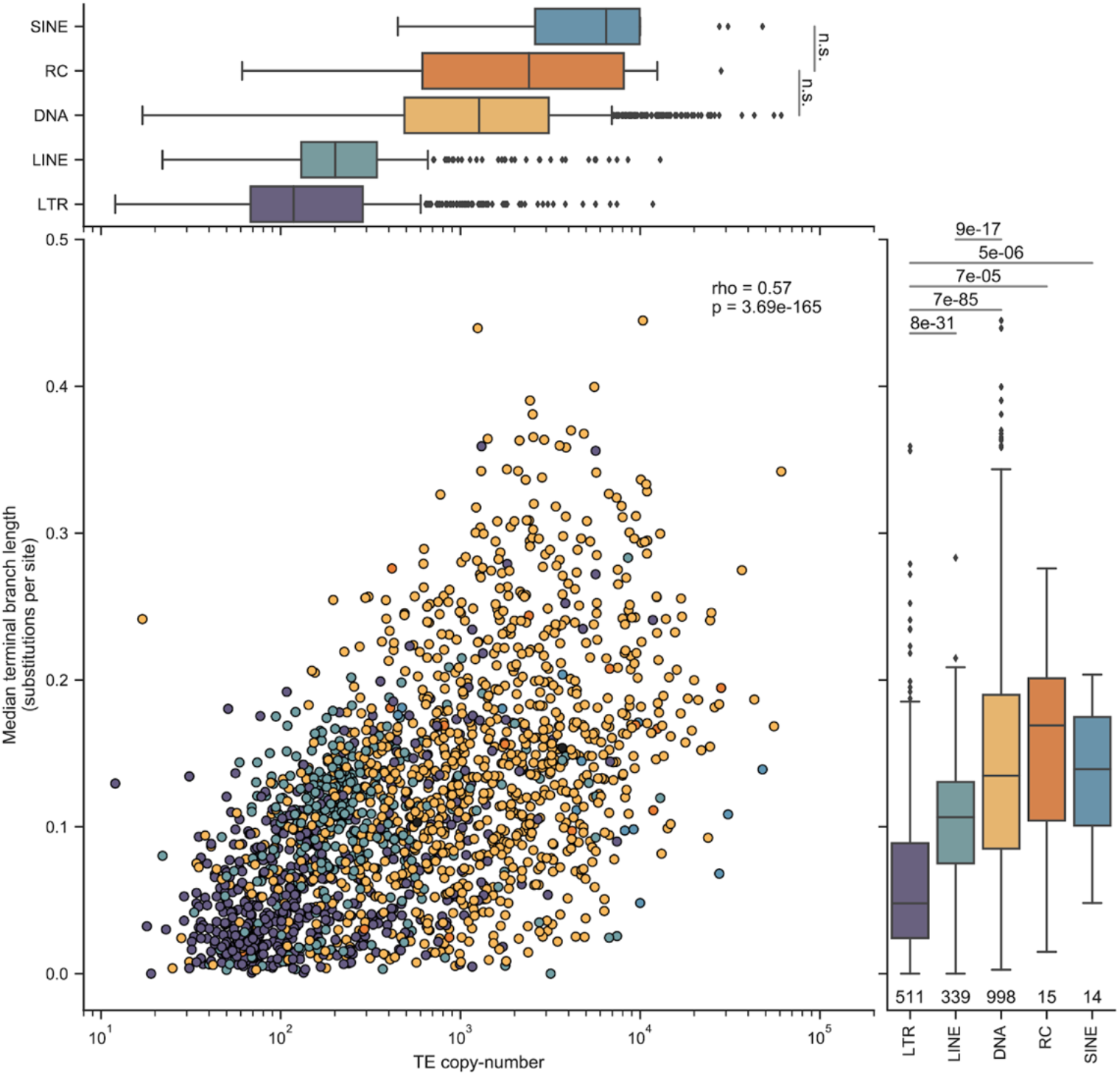
Median ages and copy numbers differ between TE classes. *Overall, there is a moderate correlation between copy number and age (Spearman’s* ρ*=0.57, p=3.69e-165). LTR elements on average are younger than other classes (lower values on the y-axis), and DNA transposons are typically older. Numbers underneath the box plots are the number of distinct TE families used in this analysis. Significance was calculated using Wilcoxon rank-sum tests between each TE class, using a threshold Bonferroni-corrected p-value of 0.001 for determining significance. For clarity, only the two non-significant tests are shown in the top panel.*

### Differential retention of TE insertions among classes

The rate at which TE insertions are removed by purifying selection is in part determined by the magnitude of their deleterious effects. Since ectopic recombination between TE copies is thought to be a major driver of selection against TEs (Blass et al., 2012; Boissinot et al., 2006; Petrov et al., 2003), increased turnover of LTRs and LINEs could be driven by selection due to their greater length relative to DNA transposons, since longer elements provide larger targets for ectopic recombination, all other factors being equal. To investigate this hypothesis, we first confirmed that there are differences in consensus sequence length between the major TE classes represented in the zebrafish genome (Supp. Fig. 2A). On average, zebrafish LTR elements are approximately 1.3 times longer than LINEs, and 4.8 times longer than DNA transposons. We then tested to see if there was a relationship between the length of TE families and their median age, and found a moderate, but significant negative correlation between the two (Spearman’s ρ = −0.35, Supp. Fig. 2B). Importantly, these correlations hold when analyzing each class separately (Supp. Fig. 2B), and thus the relationship between length and age is independent of potentially confounding differences between classes. This result is consistent with a scenario in which longer TE insertions are removed from the zebrafish genome at a faster rate than shorter insertions.

### Genomic distribution of TEs is non-random

We next looked at the distribution of TEs across chromosomes (Fig. 2A, detail 2B). Visual inspection of TE density plots reveals notable patterns in the distribution of different classes, such as the localized density peaks of RC elements (which may reflect their tendency to form tandem arrays (Pritham and Feschotte, 2007; Thomas et al., 2010)), co-enrichment of LTR elements and LINEs, and a negative correlation between LTR/LINE and SINE density. To quantify these observations, we calculated the density (as genome sequence coverage) of different TE classes in non-overlapping 2-Mb windows along the genome, then calculated the pairwise correlation between groups of interest. This approach reveals significant correlations, both positive and negative, between different TE classes (Fig. 2C). LTR and LINE density is positively correlated (Spearman’s ρ = 0.55), whereas SINE density is negatively correlated with both LINE and LTR densities (Spearman’s ρ = −0.28 and −0.45, respectively). Similar patterns of opposing LINE/SINE density have been observed in human, mouse and rat genomes, although the cause of this phenomenon is not fully understood (Gibbs et al., 2004; Lander et al., 2001; Medstrand et al., 2002; Waterston et al., 2002).

**Figure 2.**
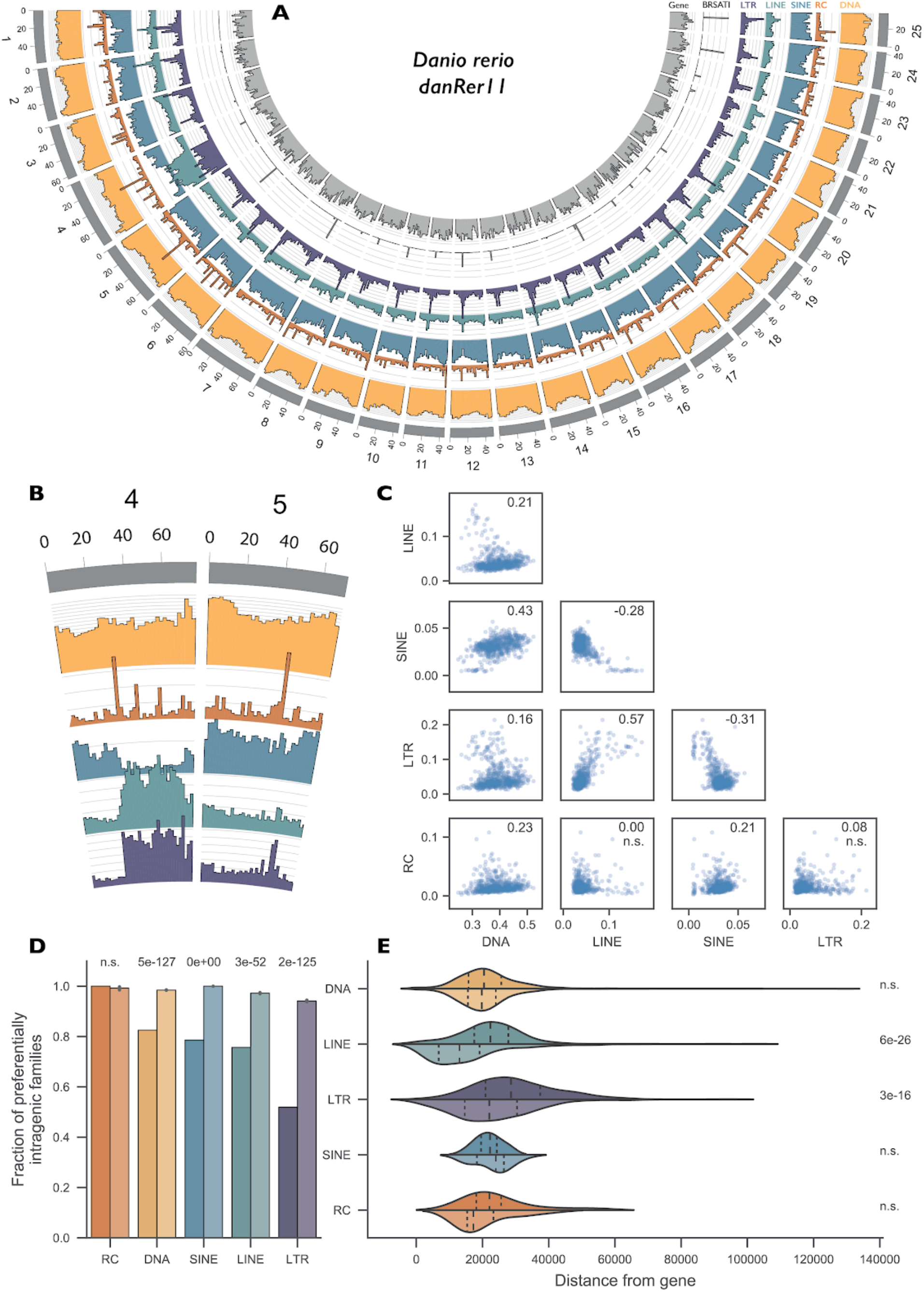
Genomic distribution of elements is non-random. *A. Genomic coverage of TEs in non-overlapping 2Mbp windows across nuclear chromosomes. Each axis line (faint grey) represents 2.5% sequence coverage. B. Detail on Chromosomes 4 and 5. C. Spearman’s rank correlations of coverage density between major TE classes. Values for* ρ *given in top-right corner of each plot; n.s. - not significant D. TE families are defined as “preferentially intragenic” if the median distance between their insertions and the closest gene is 0, i.e. most insertions in the family overlap partially or fully with gene bodies. Bars for each TE class represent observed fractions (left bars), and fractions based on random shuffling of TE insertion identities across the genome, keeping locations fixed (right bars, color desaturated). P-values calculated using binomial tests. E. Median, per family, distance of insertions from nearest genes. Top halves - distance from closest gene on same strand; bottom halves (desaturated) - distance from closest gene on opposite strand. P-values calculated using Wilcoxon rank-sum tests.*

Since LTRs and LINEs accumulate in one particularly dense cluster within each chromosome (Fig. 2A), we reasoned that these could correspond to pericentromeric regions. To corroborate this idea, we compared the density of the satellite repeat BRSATI, a marker of pericentromeric DNA (Howe et al., 2013; Phillips and Reed, 2000) to that of LTR and LINE. We found that both LTR and LINE densities were positively correlated with BRSATI density (Spearman’s ρ = 0.26, p = 5.3e-12 and ρ = 0.20, p = 7.2e-8 for LTRs and LINEs respectively). Thus, LTR and LINE achieve their highest density in pericentromeric regions. Also of note is the striking enrichment of retroelements on the long arm of chromosome four (4q), as previously observed (Howe et al., 2013). Since this region is large and thus may be driving some of the observed correlations between the density of TE classes, we repeated the analyses with chromosome 4 omitted, but observed no substantive changes in effect size or significance.

Patterns in chromosomal TE distributions are shaped both by insertion site preference of the TEs and by natural selection acting after insertion to differentially retain elements inserted in various genomic locations. To disentangle these effects, we regenerated the circos density plot shown in Fig. 2A using only insertions less than 1% diverged from their family consensus sequence (i.e., young), and those more than 15% diverged (i.e., old) (Supp. Fig. 3). Looking at the distribution of young insertions, we see not only that there is still an abundance of LTR and LINE elements in pericentromeric regions and chromosome 4q, but that the density of young DNA element insertions is also much higher on chromosome 4q than elsewhere in the genome. In contrast, older insertions of any class are depleted on chromosome 4q (Supp. Fig 3B). These results suggest that the enrichment of TEs on chromosome 4q may reflect preferential insertion of TEs on this chromosome arm and/or the fact that TEs turn over more rapidly on this arm than elsewhere in the genome.

We next examined the distribution of TE families relative to genes (Fig. 2D). The zebrafish genome is relatively gene-dense, with approximately 60% of the chromosomal DNA comprising genic regions (~3% for protein coding sequence), as defined by full-length Ensembl gene annotations on assembled chromosomes. Thus, in the absence of insertion site preference or selection, we would expect the majority of TEs to overlap genic regions. To test whether or not differences exist in the retention of different TE classes across different genic compartments, we categorized each TE family as being either preferentially intragenic if more than 50% of its copies overlapped with gene bodies, or preferentially intergenic otherwise. Then, for each TE class, we calculated the fraction of TE families categorized as preferentially intragenic and compared this to the fraction based on random shuffling of TE identities. With the exception of RC families (n=16), all families were significantly less likely to be preferentially intragenic than expected, consistent with selection against insertion within genes (Fig. 2D).

To further investigate the distribution of TEs relative to genes, we looked in more detail at intergenic insertions. For each family, we measured the median distance of intergenic insertions to the nearest gene on the same strand, and on the opposite strand (Fig. 2E); we find that LTR elements and LINEs are located significantly further away from genes on the same strand than those on the opposite strand (Wilcoxon rank-sum tests: p=3e-16 and p=6e-26, respectively). Autonomous retroelements often encode strong cis-regulatory sequences capable of affecting nearby gene expression, including promoters, splice sites, and polyadenylation signals (Clayton et al., 2020; Ishiuchi et al., 2015; Ng et al., 2020). Thus, there may be stronger selection against zebrafish LTR elements and LINEs when they insert on the same strand as a nearby gene, similar to what has been observed in mammalian genomes (Medstrand et al., 2002).

### Stage-specific regulation of TEs during early development

To investigate TE expression during zebrafish development, we took advantage of a publicly available RNA-seq dataset covering 18 stages from 1-cell to 5 days post fertilization (White et al., 2017). This high-quality poly-A pull-down stranded dataset, with five biological replicates per time-point, constitutes an ideal resource to examine gene and TE expression during early development at a high temporal resolution. In order to evaluate TE expression, we benefitted from the recent development of computational tools that allowed us to analyze expression of individual TE loci. To do so, we used STAR (Dobin et al., 2013) to map RNA-seq reads to the genome and Telescope (Bendall et al., 2019) to quantify the amount of reads mapping to individual TE copies.

TEs are abundant throughout the genome and can be incorporated into gene transcripts, for example through integrations overlapping coding sequences and UTRs (Attig et al., 2019; Kelley and Rinn, 2012; Wang et al., 2014), or as a result of intron retention events (Zaghlool et al., 2013). In such cases, it can be challenging to determine if a TE-mapping read originates from a gene promoter or a TE promoter (Lanciano and Cristofari, 2020). To address this issue, we categorized the TE annotation based on the TE position with respect to genes (Fig. 3A). Reads mapping to TEs overlapping annotated exons, UTRs, or introns of expressed genes in the same orientation were considered as transcribed in a gene-dependent manner. These TE-containing transcripts are likely to originate from the host gene’s promoter. Conversely, reads mapping to intergenic TEs or TEs in introns of genes that were not detected as expressed in any sample were considered to be driven by their own promoter, or self-expressed (Fig. 3A, Methods). Self-expressed TEs (LTR and LINE in particular) were found to be generally younger than gene-dependent TEs (Fig. 3B, Supp. Fig. 4A), which may reflect their greater likelihood to retain active promoters (Chuong et al., 2017).

**Figure 3.**
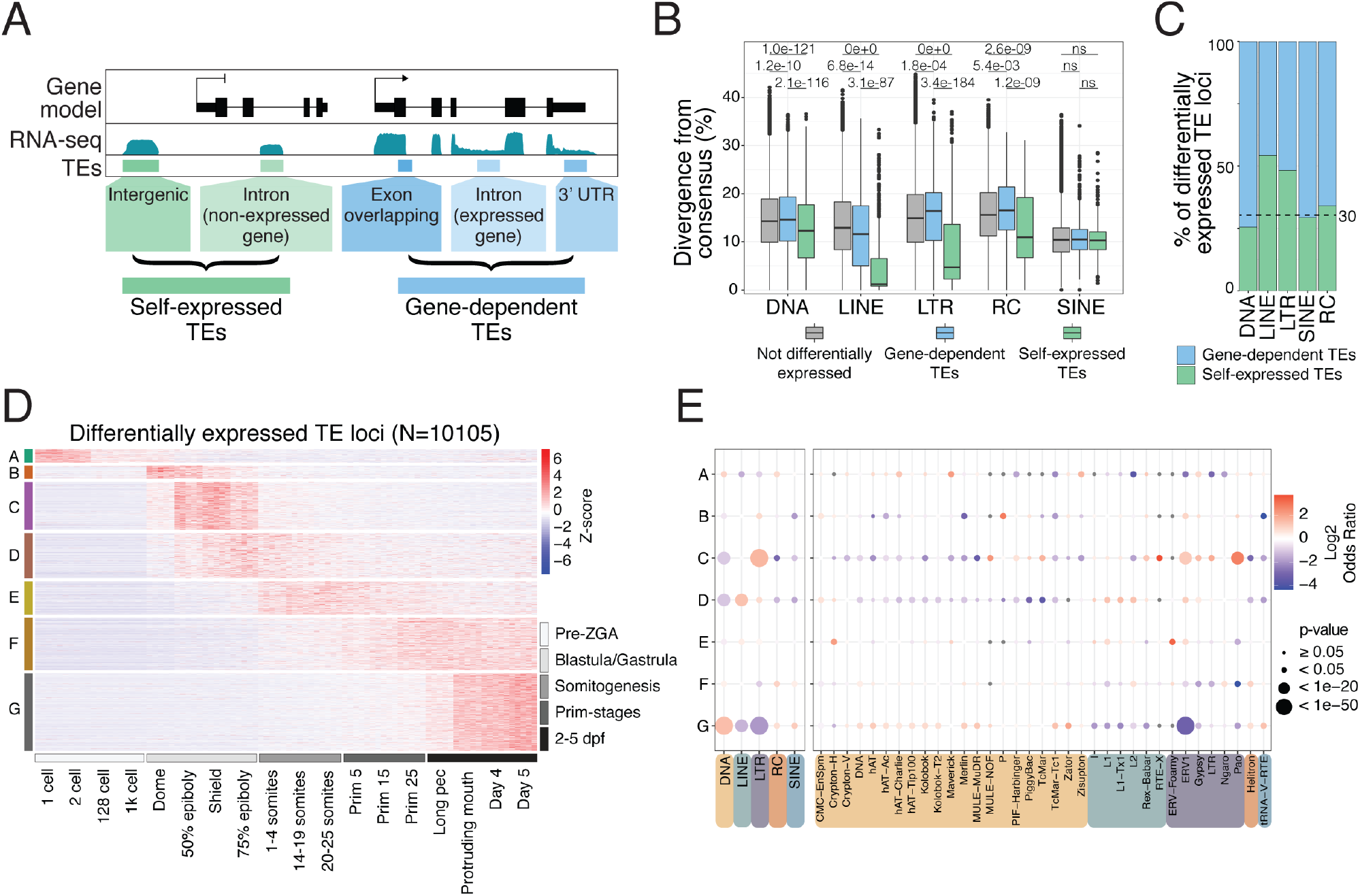
TEs are expressed in stage specific patterns during zebrafish development. A. Schematic representation of self-expression or gene-dependent expression of TE loci. B. TEs that are both differentially expressed and self-expressed are younger, with lower divergence from consensus, compared to differentially expressed gene-dependent TEs and non-differentially expressed TEs (see Supp. Fig. 4A to see the divergence from consensus for all TE categories shown in panel A). C. Fraction of differentially expressed TE loci gene-dependent or self-expressed split by TE class (see Supp. Fig. 4C for split by TE family). D. Z-score from whole embryo RNA-seq data (White et al. 2017) shows a subset of differentially self-expressed TE loci displaying stage-specific expression. Clusters are derived using k-means clustering. E. TE class-specific (left) and superfamily-specific (right) enrichment analysis per expression cluster in D. Only TE superfamilies with significant enrichment are shown. Grey dots = not significant.

Additionally, we observed a high number of alternative transcription termination sites that were not annotated in the reference transcriptome (GRCz11.98) (Supp. Fig. 5A). To prevent a TE embedded within these extended 3’ UTRs from being categorized as self-expressed, we used a transcript assembly strategy to capture all the extended 3’ UTRs (see Methods). TEs overlapping these extended 3’ UTRs were considered gene-dependent and not included for further analysis. Interestingly, our extended 3’ UTRs strongly coincide with revised gene annotations recently reported by Lawson et al., 2020 (Supp. Fig. 5B-C). Overall, we determined that self-expressed TE-derived reads account for around 0.6% and 2.5% of the reads in pre-ZGA and post-ZGA stages respectively (Supp. Fig. 4D). Together, these filtering strategies ensure that the subsequent TE differential expression analysis highlights changes in expression derived from the direct regulation of TEs, rather than differences in expression of their surrounding genes.

We then conducted a time course differential expression analysis to detect TEs that are transcriptionally regulated during development. To do so, we performed pairwise comparisons across all developmental stages and identified differentially expressed TEs as those with an FDR-adjusted p-value < 0.01 in any comparison. Notably, from all differentially expressed TE loci, 32% were self-expressed (Fig. 3C), while the rest were gene-dependent, highlighting the importance of differentiating these two categories. Clustering of expression profiles for self-expressed TEs revealed distinct temporal clusters, suggesting that TE expression is tightly regulated during zebrafish development (Fig. 3D). A subset of 466 TE loci (4.6%) are detectable at the zygote and 2-cell stages but are silent throughout the rest of development (Fig. 3D cluster A). Since the zebrafish embryo is transcriptionally inactive at this stage (Heyn et al., 2014), these TE transcripts are likely to be maternally deposited during oocyte maturation and subsequently degraded after zygotic genome activation (ZGA). Post ZGA, different TEs show stage-specific expression patterns spanning from the Dome stage until 5 days post fertilization (Fig. 3D). Notably, cluster B (469; 4.6%) and C (1,720; 17%) define two subsets of TE loci sharply activated post-ZGA and up-regulated during the blastula/gastrula stages. Cluster D (1,608; 15.9%) and E (1,195; 11.8%) contain TE loci which are up-regulated later in development, during somitogenesis, while clusters F (1,870; 18.5%) and G (2,777; 27.5%) mark a group of TE loci that peak in expression during later stages. Together, these data suggest that many TEs are expressed in a tightly regulated manner during zebrafish embryonic development.

Next, we performed an enrichment analysis to detect over/under-represented TE classes and superfamilies within each TE expression cluster (Fig. 3E). DNA transposons were generally enriched in clusters with late (larval) expression and depleted in clusters corresponding to the blastula and gastrula stages. By contrast to DNA transposons, retroelements – and LTR elements in particular – were generally enriched in clusters marking earlier developmental stages (Fig. 3E). Specifically, the LTR superfamilies ERV1, Gypsy and Pao were enriched within cluster C, which marks early post-ZGA expression at the blastula/gastrula stages. Most LINE superfamilies (I, L1, Tx1 and L2) were enriched within cluster D, which corresponds to the late stage of gastrulation and early somitogenesis (Fig. 3E). Together, these results suggest that different TE classes and superfamilies have distinct expression profiles during zebrafish development, including a pronounced activation of LTR retroelements shortly after ZGA, when early cell fate decisions are made.

### Single-cell RNA-seq resolves somatic TE expression during early embryogenesis

Our analysis of whole-embryo RNA-seq data suggests that many zebrafish TE families display stage-specific expression patterns during embryonic development. To investigate cell type- and lineage-specific TE expression during early development, we turned to a publicly available single-cell RNA sequencing (scRNA-seq) dataset (Farrell et al., 2018). This dataset spans 12 developmental stages, ranging from 3.33 hours post fertilization (hpf, so-called high stage) to 12 hpf (6-somite stage), allowing us to track TE expression along specific developmental trajectories. We re-aligned scRNA-seq reads to the most recent zebrafish genome assembly (GRCz11) and annotated reads to both genes and self-expressed TE loci defined from our bulk RNA-seq analysis. After data processing, we excluded potential cell doublets and cells with low complexity transcriptomes and high proportions of mitochondria RNAs (Farrell et al., 2018). We were left with data spanning 44,709 cells across all stages. Approximately 2% of reads mapped to self-expressed TE loci, which is comparable with what we observed with the whole-embryo RNA-seq data. Because of the shallow sequencing depth of scRNA-seq and the repetitiveness of TEs, it is difficult to confidently assess expression at individual TE loci (He et al., 2021; Shao and Wang, 2021). Thus, we analyzed the expression profile of TE at the family level by counting all reads mapping to loci from the same TE family.

To identify TE families expressed in specific cell types, we grouped all cells across the 12 stages into 68 cell-type clusters, based on both gene and TE family expression (see methods). To validate these clusters, we verified that they had captured known marker genes for distinct cell lineages, such as primordial germ cells (PGCs), the enveloping layer and notochord cells (Farrell et al., 2018, Supp. Fig. 6A), and correctly separated cells based on their developmental stages (Supp. Fig. 6B). To identify differentially expressed TE families between cell clusters, we compared TE expression levels within each cluster to their expression levels in the rest of the cells. To avoid possible technical noise from scRNA-seq (Farrell et al., 2018; Shao and Wang, 2021), we focused on TE families that are expressed in more than 10% of cells in at least one cell cluster, using only reads mapping to the self-expressed loci identified from bulk RNA-seq. Among those, 42 TE families were significantly upregulated in at least one cell cluster compared to all other cells (Fig. 4A). Using hierarchical clustering analysis, these families can be divided into two broad categories: a group of 14 TE families that are highly expressed in the blastula (largely undifferentiated cells) and gastrula stage but not later in development, and the other group of 28 TE families that are expressed much later in development, when most cells have already differentiated into distinct cell lineages. In agreement with our bulk RNA-seq results for retrotransposons, the first group (early expression) includes members of the Gypsy, ERV and L1 superfamilies and a few members of the EnSpm (CMC), Helitron, P-element, hAT and Harbinger superfamilies of DNA transposons. The second group (late expression) includes representatives from all the retrotransposon superfamilies identified in the first group, but only EnSpm and Helitron DNA transposons. We found no significant differences in age or TE classes between those two broad groups of elements (Mann-Whitney U test, p=0.66). Overall, TE families belonging to the endogenous retrovirus ERV superfamily were strongly enriched across both groups (16 out of 42 TE families), compared to their representation in the genome (54 out of 1931 TE families, Fisher’s exact test, p < 0.001).

**Figure 4.**
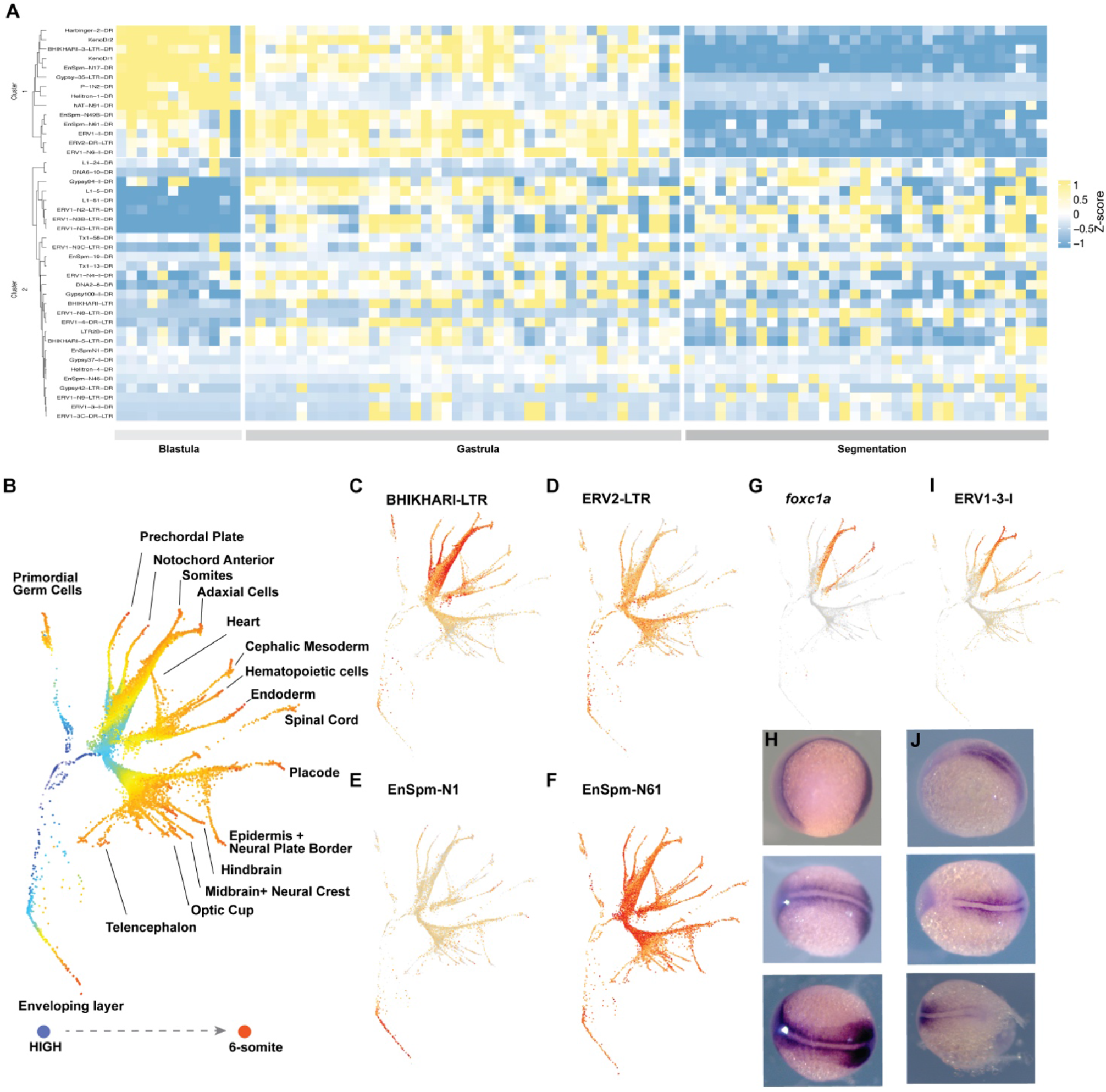
TE families with cell-lineage expression across development stages. A. Heatmap of differentially expressed TE families between cell clusters across developmental stages. Hierarchical clustering shows two groups of TE families with distinct expression patterns-one group with early expression in blastula and gastrula stages and one group with later expression in gastrula and segmentation stages. TE classes are equally represented in both groups. B. Pseudotime tree across 12 development stages based on both gene and TE expression. C-F. TE families with expression patterns in different cell lineages. C. BHIKHARI-LTR, D. ERV2-LTR, E. EnSpm-N1, F. EnSpm-N61. G. The expression pattern of foxc1a in pseudotime tree and H. in 11 hpf embryos by in situ hybridization. I. The expression pattern of ERV1-3-I in the pseudotime tree and J. in 11 hpf embryos by in situ hybridization.

To visualize TE families with late expression patterns along developmental trajectories, we conducted a pseudotime tree analysis (Farrell et al., 2018, Fig. 4B). Intriguingly, we found that several TE families were highly expressed in specific somatic cell lineages (Fig. 4D-F). One of the most striking trajectories was that of BHIKHARI, an ERV1 family expressed exclusively in the mesendoerm and in PGCs (Fig. 4C). These results corroborate earlier reports that BHIKHARI transcripts marks specifically the developing mesendoderm of zebrafish (Vogel and Gerster, 1999). Importantly, using uniquely-mapping reads we could infer that BHIKHARI expression is driven by the majority of BHIKHARI copies (97 out of 98 self-expressed BHIKHARI loci) dispersed throughout the genome, and not a single or a few isolated copies (Supp. Fig. 7A). Another striking example was ERV1-3, which is highly expressed in axial and paraxial mesoderm after 50% epiboly stage (Fig. 4I). Again, ERV1-3 expression was driven by multiple insertions in the genome (71% of reads were from 10 out of 45 self-expressed loci; Supp. Fig. 7B), suggesting that this expression pattern is driven at least in part by ERV1-3’s own promoter activity rather than the local genomic environment. To experimentally validate these observations, we conducted *in situ* RNA hybridization using a probe designed against the *pol* gene of ERV1-3 on embryos at the 3-somites stage (11 hpf). As a comparison, we also performed *in situ* hybridization with a probe for *foxc1a*, a transcription factor known to mark the paraxial mesoderm (Topczewska, 2001; Wilm et al., 2004, Fig. 4G). The results show that ERV1-3 and *foxc1a* RNA transcripts have very similar expression patterns in zebrafish embryos and both specifically mark the paraxial mesoderm (Fig. 4H-J).

Although transcripts of DNA transposons are mostly driven by nearby genes (Fig. 3C), several families are self- and differentially expressed between cell clusters. Among those, members of the EnSpm superfamily were enriched in our analysis (Fisher’s exact test, p=0.025). Most of the EnSpm families with lineage-specific expression are nonautonomous elements with no detectable coding sequences. Yet, analysis of uniquely mapping reads indicates that the expression of each EnSpm family was driven by multiple loci throughout the genome. For instance, 11% (202/1898 loci) of EnSpm-N61 copies showed evidence of expression in most cell lineages during gastrulation (Fig. 4F). Together these data suggest that diverse TE families encompassing both retrotransposons and DNA transposons display specific pattern of spatiotemporal expression in developing zebrafish embryos.

## Discussion

In this work we have carried out a comprehensive analysis of the zebrafish TE ecosystem and their embryonic niche using a wealth of transcriptomic data spanning developmental stages from pre-ZGA to five days post-fertilization. The zebrafish genome contains nearly 2,000 TE families from all major classes and superfamilies, approximately 65% of which are expressed during development. From analyses of both bulk and single-cell expression data, our results suggest that zebrafish TEs span a wide diversity of expression patterns, from highly stage- and cell type-specific expression, to broad expression throughout development. These patterns vary both between TE classes and within superfamilies and are in part reflected in the broad differences in their genomic distribution.

Measuring expression of TEs remains a challenge in genomic analyses due to their repetitive nature, intricate transcriptional relationship with host gene expression, and the general complexity of the transcriptome (Lanciano and Cristofari, 2020). Short reads mapping to TE sequences cannot easily distinguish whether they derive from a TE promoter or are part of a gene or readthrough transcript of sorts, including non-coding RNAs which are ubiquitous in vertebrate genomes (Kung et al., 2013) and often contain TEs (Kapusta et al., 2013). Recent studies have attempted to address these technical difficulties by analyzing exon-overlapping, intronic and intergenic reads separately, both with bulk and scRNA-seq data (He et al., 2021; Kong et al., 2019; Shao and Wang, 2021). In this work, we have combined Telescope, a recently developed tool to detect TE expression at single loci resolution from bulk RNA-seq data (Bendall et al., 2019), with genome-based classification to differentiate between TE expression most likely derived from gene promoters or from TE promoters (Fig. 3A, see methods). This approach suggests that around two thirds of TE-mapping reads in the zebrafish transcriptome are most likely associated with host gene expression and readthrough transcription. Thus, the majority of TE sequences in zebrafish are not expressed from their own promoters, but are expressed as part of chimeric read-through transcripts, both coding and non-coding.

TE fragments embedded in gene transcript isoforms may have diverse functional consequences. For example, they have been shown to be the source of new protein coding-exons, RNA binding motifs and microRNA target sites (Cosby et al., 2021; Lev-Maor et al., 2003; Petri et al., 2019; Zarnack et al., 2013). The high fraction of TE fragments co-transcribed with host sequences revealed by our analysis beg for further investigation of their functional significance in zebrafish development. An interesting case is the Maverick/Polinton class of DNA transposons, which is strongly enriched at zygotic and pre-ZGA stages. Maverick/Polinton elements have been associated with the DNA 6mA modification during early embryonic zebrafish development, hinting at the unusual regulation of this family of TEs at this stage (Liu et al., 2016). It is also important to emphasize that our definition of self-expressed TEs is conservative and may underestimate the activity of TE-derived promoters. For example, we noticed that among differentially expressed gene-dependent TE loci, LTRs were the TE class with the highest fraction of overlap with 5’ UTR and coding exons (Supp. Fig. 4B). These may represent chimeric LTR-host gene transcripts driven by LTR rather than host gene promoters (Thompson et al., 2016). Additional analyses would be required to validate the chimeric structure and transcription start sites of these transcripts.

Using bulk and single-cell RNA-seq to untangle temporal and lineage-specific patterns of TE expression, we observe broadly distinct patterns between the major TE classes. As documented in mammalian species (Franke et al., 2017; Göke et al., 2015; Grow et al., 2015), we observe that LINE and LTR retroelement transcripts are particularly abundant at or shortly after ZGA in zebrafish. By contrast, we find that DNA transposon transcripts tend to be enriched prior to ZGA (i.e. maternally deposited) or expressed later in development. We also note that retroelement insertions are significantly more likely to drive their own expression than are DNA elements, which is consistent with the fact that retroelements typically encode strong promoters while DNA transposons are thought to have generally weaker or less specific promoters (Palazzo et al., 2017, 2019). This observation may also explain why we found that retroelements are less common nearby or within genes than DNA transposons, since their promoters and other cis-regulatory elements have greater potential to interfere with gene expression. This difference in genomic distribution may also be partly driven by the length of the elements, which is likely correlated with the frequency at which they contribute to ectopic recombination (Petrov et al., 2011). Indeed, LINEs and LTR retroelements are generally longer than DNA transposons. Accordingly, we found that SINEs, which are short and transcribed by RNA pol III and therefore less likely to interfere with pol II-mediated regulation, are also more closely associated with genes than other subclasses of retroelements – a trend also observed in mammalian genomes (Gibbs et al., 2004; Lander et al., 2001). Thus, the potential to interfere with gene expression and the propensity to mediate ectopic recombination likely act in concert to shape the differential accumulation of zebrafish TE classes relative to genes.

With respect to expression, certain superfamilies stand out in both the bulk and scRNA-seq analyses – most notably the ERV superfamilies. ERV elements tend to be highly expressed immediately after ZGA, often in a cell-type specific fashion and apparently using their own promoters, before being silenced later in development. This pattern suggests that ERV expression is governed by tightly regulated cis-regulatory sequences responsive to both transcriptional activators as well as repressors. This is reminiscent of mammalian ERVs which are activated by stage-specific TFs and repressed by sequence-specific KRAB-zinc finger proteins (Bruno et al., 2019; Hermant and Torres-Padilla, 2021). Given the lack of the KRAB domain in zebrafish, a clear research avenue for the future will be to identify the transcriptional regulators silencing zebrafish ERVs. Compared with other TE types, ERVs appear to be more intimately tied up in the host embryonic development process, and this raises the possibility that they are able to influence embryogenesis to an extent that we have not yet fully appreciated.

Another intriguing finding of our study is the identification of a small subset of TE families (e.g. BHIKHARI and ERV1-3, from the aforementioned ERV superfamily, and EnSpm-N61, a DNA transposon) with high level of RNA expression in somatic progenitor cell lineages. Could such somatic expression facilitate transposition in the germline? One possibility is that somatic expression provides an indirect route for TEs to enter the germline; this has been observed during oogenesis in *Drosophila melanogaster*, where TEs expressed in support cells surrounding the oocytes either infect, or are trafficked into, mature oocytes (Chalvet et al., 1999; Wang et al., 2018). Alternatively, it may be that somatic expression is only mildly deleterious to the zebrafish host, and therefore occasionally arises with little functional consequence for either the host or the TEs. This may be the case if the resulting protein products are non-toxic and if somatic transposition events remain rare. Finally, it is possible that spatiotemporal patterns of TE expression may occasionally support organismal development. Whilst this is a provocative idea, there are now several of examples of TEs with important roles in embryonic development: L1 and MERVL in mice (Jachowicz et al., 2017; Macfarlan et al., 2012; Percharde et al., 2018), HERV-K and HERV-H in humans (Grow et al., 2015; Lu et al., 2014), and ERNI in chicken (Blanc et al., 2014); the last of these is noteworthy as it functions in a strictly somatic niche. Functional experiments will be necessary to determine whether zebrafish TEs expressed in somatic lineages reflect selfish, neutral, or mutualistic behaviors, and we anticipate that this will be a fruitful topic of study in coming years.

The activity of retroelements during early embryonic development has been noted in many vertebrate species, particularly in mammals. Many of the patterns observed in these studies are recapitulated in zebrafish, indicating that features such as robust expression following ZGA, lineage-specific expression and accumulation in gene-poor regions, are features shared by diverse retroelement superfamilies across a broad range of vertebrates. In contrast, much less is known about the behavior of DNA transposons during development, largely due to the paucity of active DNA transposon families in mammals, with the notable exception of vespertilionid bats (Platt et al., 2016). Unlike LINEs and LTR elements, DNA transposon-derived transcripts are enriched both very early in development (prior to ZGA) and in the latest stages of development (4-5 dpf). Mechanisms to prevent activation and mobilization of TEs in germ cells, such as Piwi-interacting RNA (piRNA) pathway, have been described in zebrafish (Houwing et al., 2007, 2008; Kaaij et al., 2013). piRNAs and Piwi proteins are maternally deposited and localized in the germ plasm (Houwing et al., 2007). Following the first cell divisions, cells that inherit the germ plasm will develop into PGCs (Raz, 2003). Zebrafish piRNAs are enriched in LTR targets and contain fewer DNA transposon targets, indicating a greater degree of protection against younger LTR elements compared to DNA transposons (Houwing et al., 2007; Kaaij et al., 2013). Thus, the depletion of LTR transcription in pre-ZGA stages, which mainly contain maternally deposited transcripts, may be due to efficient repression by the piRNA pathway. Interestingly, piRNAs targeting Harbinger DNA transposons are abundant in zebrafish ovaries, explaining the depletion of this superfamily at early stages (Fig. 3D, Houwing et al., 2007). Our analyses reveal that DNA transposon transcription is more often gene-dependent than retroelement transcription. One feature of DNA transposons that may facilitate their hijacking of host promoters is the presence of splice sites within their sequence, which has been implicated in the formation of chimeric transcripts which occasionally encode transposase-host fusion proteins coopted for cellular function (Cordaux et al., 2006; Cosby et al., 2021; Newman et al., 2008). This raises the question of what proportion of zebrafish DNA transposon transcripts are capable of producing chimeric protein products.

With its remarkably rich TE content, zebrafish, more than most, exemplifies the idea of the genome as an ecosystem. Much like the species they parasitize, TEs possess traits that are shared across taxonomic groups, but also traits that are unique to each family. For almost all TEs however, embryonic development is a critical period for their long-term success, and increasingly it is clear that many TEs are not idle passengers in the process. Zebrafish is a powerful model for the study of vertebrate embryogenesis, and yet is only beginning to attract interest as a system for studying genome evolution and the role of TEs during the process. We hope that this work provides a useful foundation for the development of zebrafish as a model for investigating the fascinating interplay between TEs and their hosts.

## Author contributions

J.W. performed genomic TE analysis. Q.R. performed bulk RNA-seq analysis. N.C. performed scRNA-seq analysis. C.F. and J.M.V supervised the data analysis and interpretation. All authors discussed the results and contributed to writing the final manuscript.

## Acknowledgments

This work was supported by grant R35-GM122550 from the National Institutes of Health to C.F. Work in the Vaquerizas laboratory is supported by the Max Planck Society, the Deutsche Forschungsgemeinschaft (DFG) Priority Programme SPP2202 ‘Spatial Genome Architecture in Development and Disease’ (project number 422857230 to J.M.V.), the DFG Clinical Research Unit CRU326 ‘Male Germ Cells: from Genes to Function’ (project number 329621271 to J.M.V.), the European Union’s Horizon 2020 research and innovation program under the Marie Skłodowska-Curie (grant agreement 643062, ZENCODE-ITN to J.M.V.), the Medical Research Council, UK (award reference MC_UP_1605/10 to J.M.V.), the Academy of Medical Sciences and the Department of Business, Energy and Industrial Strategy (award reference APR3\1017 to J.M.V.). J.N.W is supported by a Human Frontier Science Program long-term fellowship (LT000017/2019-L). N.C is supported by a Distinguished Scholar Award from the Cornell Center for Vertebrate Genomics. We would like to thank Dr. Joseph Fetcho for zebrafish husbandry support. Finally, we would like to thank members of the Vaquerizas and Feschotte labs for valuable feedback and discussion.

## Methods

### Transposable element annotation

TEs were mapped to the zebrafish genome (May 2017, GRCz11/danRer11, accessed from the UCSC genome browser) using RepeatMasker version 4.08 (Smit et al., 2013, http://www.repeatmasker.org). For mapping, we used the rmblastn engine (version 2.2.27+) and the Dfam_Consensus-20181026 and RepBase-20181026 libraries. The following parameters were set: -a, -s, -nolow, -gccalc, -gff, -cutoff 200, -no_is. The RepeatMasker output files were processed using ParseRM (Kapusta et al., 2017; https://github.com/4ureliek/Parsing-RepeatMasker-Outputs); this was used to generate measurements of Kimura CpG-corrected percent-divergence from consensus sequence. TE copy number estimates were acquired from the output of the perl script onecodetofindthemall.pl (Bailly-Bechet et al., 2014), which reconstructs fragmented repeats and full-length LTR elements.

### Dating TE insertions

To build phylogenetic trees for each TE family, defragmented sequences were extracted from the genome and aligned using MAFFT v7.419 (Katoh and Standley, 2013), with the --auto flag set to true. A minimum sequence length of 100 was specified for inclusion in the alignments. TEs with fewer than 10 suitable sequences were ignored, and alignments of TEs with high copy number were restricted to a random selection of 1,250 sequences, in order to enable computation in a reasonable time frame. FastTree v2.1.10 (Price et al., 2010) was used to construct approximate maximum-likelihood phylogenetic trees, using a generalized time-reversible model. Branch lengths were rescaled to optimize Gamma20 likelihoods. For a given TE insertion, the age was specified as the branch length from the leaf to the most recent ancestor (terminal branch length); for a family of TEs, the average age was calculated as the median of the terminal branch lengths. RepeatMasker-derived divergence from consensus sequence was used as an alternative measure of age in Supp. Fig. 3.

### Analysis of genomic distribution

To visualize the genomic distribution of different TE classes, we split the genome into 2Mbp windows and calculated TE coverage as the percentage of TE-derived base pairs in each window. The results were plotted with Circos (Fig. 2A, Krzywinski et al., 2009). The relationship between the distributions of different TE classes were calculated with Spearman’s rank correlation on the windows, sorted by percent-coverage.

To investigate the distribution of TEs relative to genes, the distance between each insertion and the nearest gene was measured using bedtools (Quinlan and Hall, 2010). For a given TE family, if the median distance between insertions and genes was equal to zero, we described that family as “preferentially intragenic”. We then compared the observed fraction of preferentially intragenic families to that expected based on random shuffling of TE labels throughout the genome, thus keeping the overall TE distribution the same but removing differences between families. The significance of the difference between observed and expected intragenic fractions for each TE class was assessed using binomial tests. Lastly, for each family we compared the median distance of insertions to genes on the same strand and different strands. We then compared the distribution of these estimates for each TE class, comparing distance on same strand and different strand using Wilcoxon rank-sum tests.

### TE loci classification

ChIPseeker (Yu et al., 2015) was used to annotate the TEs position with respect to protein coding genes (GRCz11 annotations, release 98) with the following genomic priority: genomicAnnotationPriority = c(“5UTR”, “3UTR”, “Exon”, “Promoter”, “Intron”, “Downstream”, “Intergenic”). If a TE was annotated as exonic or in 5’ or 3’ UTR region, it was considered “Exon overlapping”. For intronic TEs, if the residing gene had more than 10 normalized counts in at least 1 sample, the TE was considered “Intron expressed gene”. On the other hand, if an intronic TE was within a gene that did not had at least 10 normalized counts at any sample, then it was considered “Intron non-expressed gene”. TEs overlapping extended 3’ UTR regions (see details below) were considered “Extended 3’ UTR”. The rest of TEs were considered “Intergenic”. “Exon overlapping”, “Intron expressed gene” and “Extended 3’ UTR” were considered gene-dependent TEs. On the other hand, “Intron non-expressed gene” and “Intergenic” TEs were considered self-expressed. TEs fragments reconstruct as part of the same TE by onecodetofindthemall.pl were given the same TE classification in the following hierarchy: Exon > Extended 3’ UTR > Intron expressed gene > Intron non-expressed gene > Intergenic.

### Bulk RNA-seq mapping

RNA-seq data (White et al., 2017) was downloaded from European Nucleotide Archive accession ERP014517. Paired end reads were trimmed using BBduk (http://jgi.doe.gov/data-and-tools/bbtools) with the following parameters: ktrim=r, k=23, mink=11, hdist=1, tbo, tpe. Trimmed read were mapped to the GRCz11 zebrafish genome with appended ERCC spike-in sequences using STAR (version 2.5.2b, Dobin et al., 2013) with the following parameters: --chimSegmentMin 10 --winAnchorMultimapNmax 200 --outFilterMultimapNmax 100. STAR genome index was generated giving GRCz11.98 ensemble annotations with the parameter –sjdbGTFfile. Alignment files were sorted and indexed using sambamba (version 0.6.7, Tarasov et al., 2015). TEtranscripts (Jin et al., 2015) was run to obtain gene counts with the following parameters --stranded reverse --mode multi. Telescope (Bendall et al., 2019)was used to obtain TE counts at TE loci resolution. Since telescope does not consider stranded RNA-seq data, alignment files were split between forward and reverse mapping strand using ad hoc script with samtools (Li et al., 2009, version 1.10) to subset based on sam flags. Forward alignment files were counted to forward orientation TEs and reverse alignment files were counted to reverse orientation TEs using telescope assign (version 1.0.3) with the following parameters --theta_prior 200000 --max_iter 200 --updated_sam. Counts from TE fragments reconstruct by onecodetofindthemall.pl were merged.

### Bulk RNA-seq differential expression analysis

DESeq2 (Love et al., 2014, version 1.28.1) was used to perform the differential expression analysis of TE loci. ERCC spike-in mix was used to calculated a normalization factor using RUVseq (Risso et al., 2014, version 1.24.0) remove unwanted variation strategy with TEtranscripts gene counts. DESeq was run for gene and TE counts together with the following experimental design: spike-in normalization factor + developmental stage. Since gene expression accounts for a bigger fraction of the transcriptome, running DESeq on TE counts together with gene counts ensures a better dispersion estimation, that will impact DESeq normalization. To remove non-expressed genes and TEs, only genes and TEs with more than 5 reads in at least two samples were considered. To obtain TEs that were differentially expressed during development, pairwise comparisons between any developmental stage was performed. Multiple test correction for all the pairwise comparisons was performed by stacking all the result tables from each comparison in a single table and using p.adjust function with parameter method=“BH” in R (version 4.0.1) to calculate the adjusted p-value. To remove very lowly expressed signal, TEs with less than 10 reads in any stage were removed. TEs with a p-adjusted value lower than 0.01 in any pairwise comparison were considered differentially expressed. Finally, gene-dependent TE loci were discarded.

### TE loci clustering and enrichment analysis

Differentially expressed TE loci normalized counts matrix was standardized using Z-score transformation. Then, the matrix was clustered using K-mean clustering R function kmeans with the following parameters iter.max=500, nstart=50 and algorithm=“Lloyd”. After visual inspection, it was decided to limit the number of clusters to 7 since it represented most of the variance without over clustering. Heatmap representation of the matrix was produced using pheatmap function from pheatmap R package (version 1.0.12). Enrichment of TE classes and families within TE loci clusters was performed using fisher.test R function for contingency tables build by counting TEs from class X or class not X in cluster Y or not Y. Statistical p-values were corrected for multiple testing.

### Detection of extended 3’ UTR regions

StringTie (Pertea et al., 2015, version 1.3.6) was used to find the extended 3’ UTR regions not present in GRCz11.98 ensembl annotations. Alignment files from biological replicates were combined to increase sequencing depth. StringTie was run without reference annotations and with the following parameters --rf -t -c 1.5 -f 0.05. Using ad hoc R script, for each isoform of known genes the last exon was subtracted and using GRCz11.98 ensembl annotations the 3’ extended regions was calculated. Extended 3’ regions were calculated separately for each developmental stage and collapsed into a single annotation. Extraction of extended 3’ UTR regions with respect to GRCz11.98 Ensembl annotations was done similarly for data from Lawson et al., 2020 in order to compare it with this study extended 3’ UTR regions.

### Mapping and annotation of the single-cell RNA-seq

We downloaded the single-cell RNA-seq data from (Farrell et al., 2018) and re-mapped the reads to GRCz11/danRer11 using Bowtie2 (Langmead and Salzberg, 2012). We used the same parameters for Bowtie2 as described in Farrell et al., 2018. After mapping to the reference, we annotated reads to both genes and TEs using Drop-seq tools as in Drop-Seq Alignment Cookbook v2.0.0 (Macosko et al., 2015). The reference file of genes for GRCz11 was downloaded from Ensembl (GCA_000002035.4). TE reference file was created from Repeatmasker v4.08 as described, and we only annotate TE transcripts to self-expressed TE loci identified from our bulk RNA-seq analysis. After the annotation, we combined all the reads from the same TE family and only counted the expression level at TE family level. We then created a matrix of digital gene expression (DGE) for both genes and TE families using the DigitalExpression function in Drop-Seq tools (Macosko et al., 2015).

### Cell cluster identification and cell-type specific TEs

We then used the DGE matrix to filter out cells with low complex transcriptome or potential cell duplets based on total number of transcripts and genes, as described in Farrell et al., 2018. TE expression (including both gene-dependent and self-expressed) is roughly 8% of the total transcriptome, so we increased the threshold for maximum reads and genes by 8% for each developmental stage. Our thresholds for gene and unique molecular identifier (UMIs) for each development stage were: high stage (1,000-8,100 genes, 1,500-43,200 UMIs), oblong stage (625-8,100 genes, 1,500-32,400 UMIs), dome stage (800-4,104 genes, 2,000-21,600 UMIs), 30% epiboly (625-3,240 genes, 1,000-18,900 UMIs), 50% epiboly (600-4,320 genes, 1,500-27,000 UMIs), shield (600-2,700 genes, 1,000-16,200 UMIs), 60% epiboly (600-3,780 genes, 1,500-24,300 UMIs), 75% epiboly (600-3,456 genes, 1,400-21,600 UMIs), 90% epiboly (500-3,780 genes, 1,000-21,600 UMIs), bud stage (500-3,456 genes, 1,000-18,900 UMIs), 3-somite stage (500-3,240 genes, 1,000-13,500 UMIs), 6-somite stage (500-3,240 genes, 1,000-13,500 UMIs). We also excluded cells with unusually high mitochrondria content (> 45 % of total reads per cell), an indication of stressed cells or cell apoptosis. After filtering, we have 44,709 cells for downstream analysis. We then used Seurat v3.0 to correct for batch effects and identify cell clusters based on expression of both genes and TE families (*dims=114, resolution =5.6*). We then identify cell-type specific TEs based using FindAllMarkers *(min.pct = 0.1, logfc.threshold = 0.25, return.thresh=0.05)*.

### URD

To obtain the cell trajectory across the developmental stages, we constructed a pseudotime tree based on both gene and TE family expression. We use the R package *URD* to conduct a diffusion map and flood stimulation (n=1,500) for all cells, as described in Farrell et al., 2018. We then define the root as cells at high stage and tip clusters from 6-somite stage using Infomap-Jaccard clustering. We simulated 10,000 random walks for all cells between root and tip clusters, and reconstructed a cell trajectory tree from the simulation results. We then use force-directed layout to visualize the reconstructed tree.

### *In situ* hybridization

To validate the TE expression from our single-cell analysis, we performed *in situ* hybridization in zebrafish embryos as described (Thisse and Thisse, 2008). We amplified probes for ERV-2 LTR by using primers 5’-ACATNCCAGCTAGGAGGGACATT-3’ and 5’-CCTTTATTGAGACGTGTTGGTTAATCTGCAGT-3’, pol region of ERV1-3 by 5’-GATCCACAAACAGGCCAGAA-3’ and 5’-ACCTGCACACAAACATCGGA-3’, and *foxc1a* by 5’-CAGTCTTCTTGACGACTGTTCTTC-3’ and 5’-TAATCGAAATACTGGTTTGGTC-3’ from wildtype TU embryos, then cloned it into pMiniT 2.0 for *in vitro* transcription. The mRNAs of ERV2-LTR, ERV1-3-I and *foxc1a* are used as probes to hybridize embryos collected at 6.75 and 11 hpf, respectively. The RNA is labeled by DIG color and imaged by ZEISS stereo microscope.

## Supplementary Material

***Supplementary Data 1.** Primary RepeatMasker output file and summary table, along with processed data generated from ParseRM.pl (see methods). All downstream analyses and RNA-seq mapping were performed using these TE annotations.*

***Supplementary Data 2.** Multiple sequence alignments and corresponding phylogenetic trees of all families with at least 10 sequences of at least 100bp in length. See methods for construction details.*

***Supplementary Data 3.** Summary table containing genomic attributes of all TE families analyzed in this project. A README file with column descriptions is included within.*

**Supplementary Figure 1.**
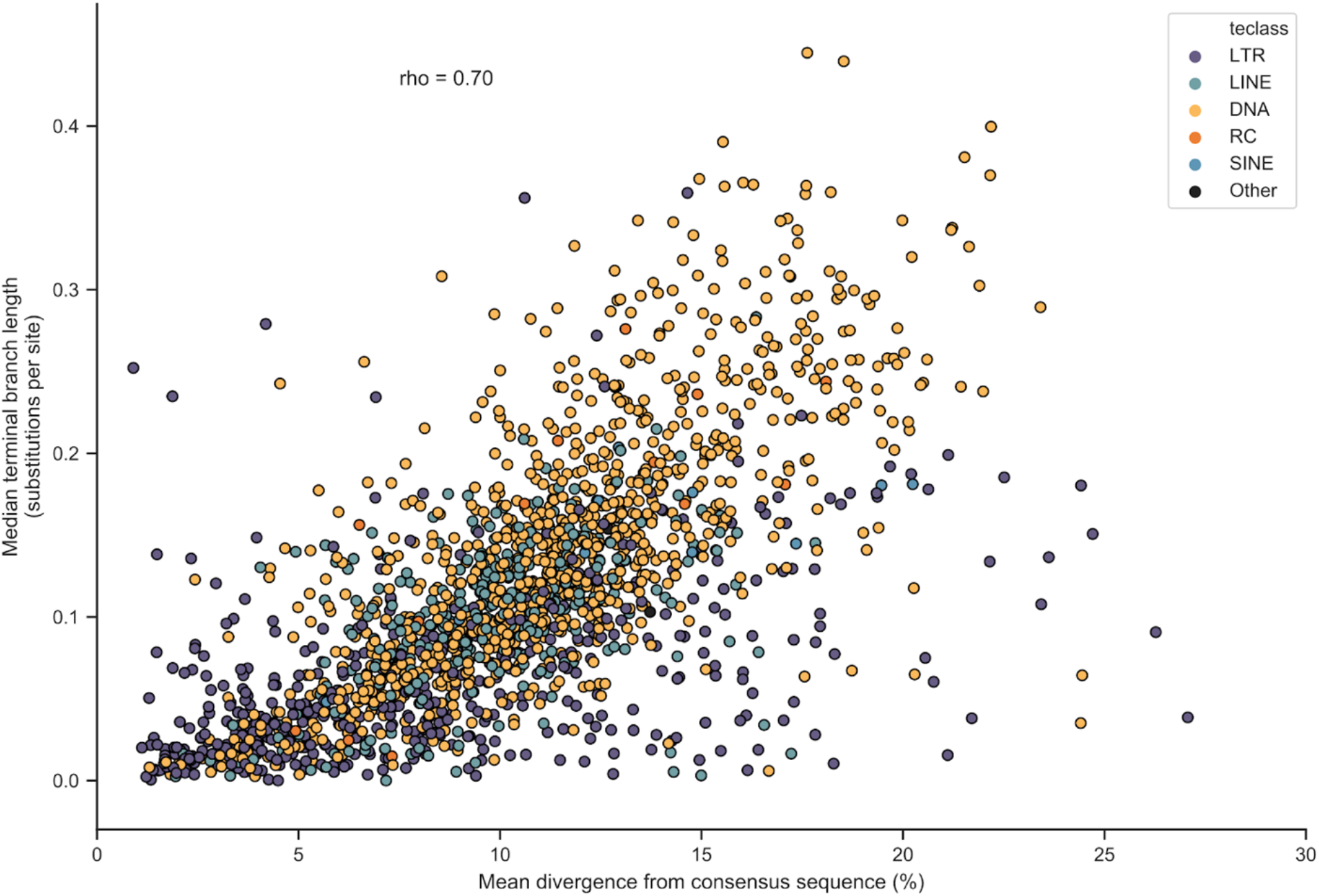
Correlation between mean divergence from consensus and median terminal branch length. This analysis serves to validate the use of terminal branch lengths as a measure of age. It correlates strongly with the more widely used metric, “divergence from consensus sequence”, and is less sensitive to inflation as a result of TE family substructure. This can be seen in the case of LTR elements, which frequently have independently active subfamilies, and for which the divergence from consensus measure increases more rapidly than median terminal branch length.

**Supplementary Figure 2.**
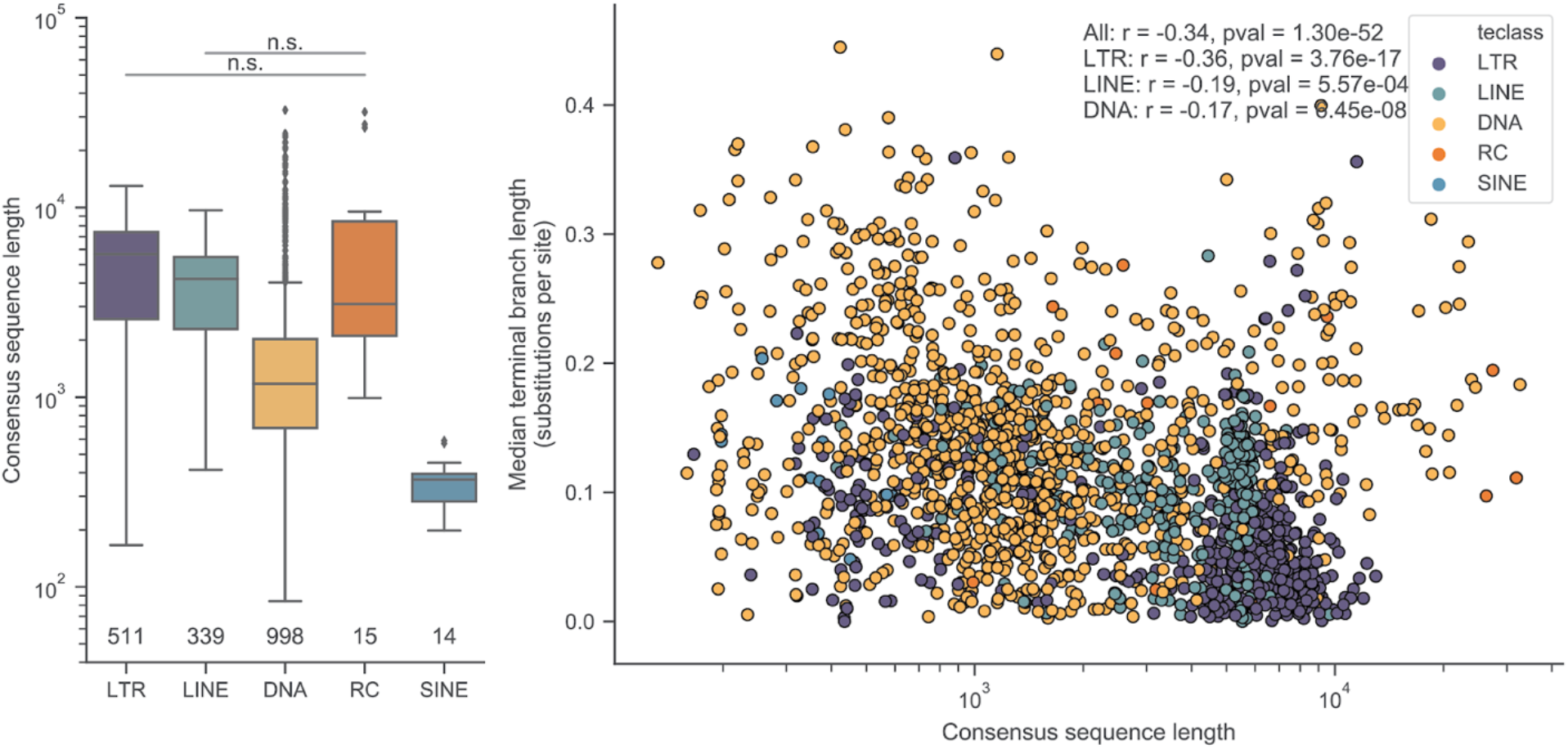
TE length correlates with age. A. Differences in consensus sequence lengths between classes. Wilcoxon rank-sum test P-values given above boxes. B. Correlation between consensus sequence length and age for different TE classes. Values for r are calculated with Spearman’s rank correlation test.

**Supplementary Figure 3.**
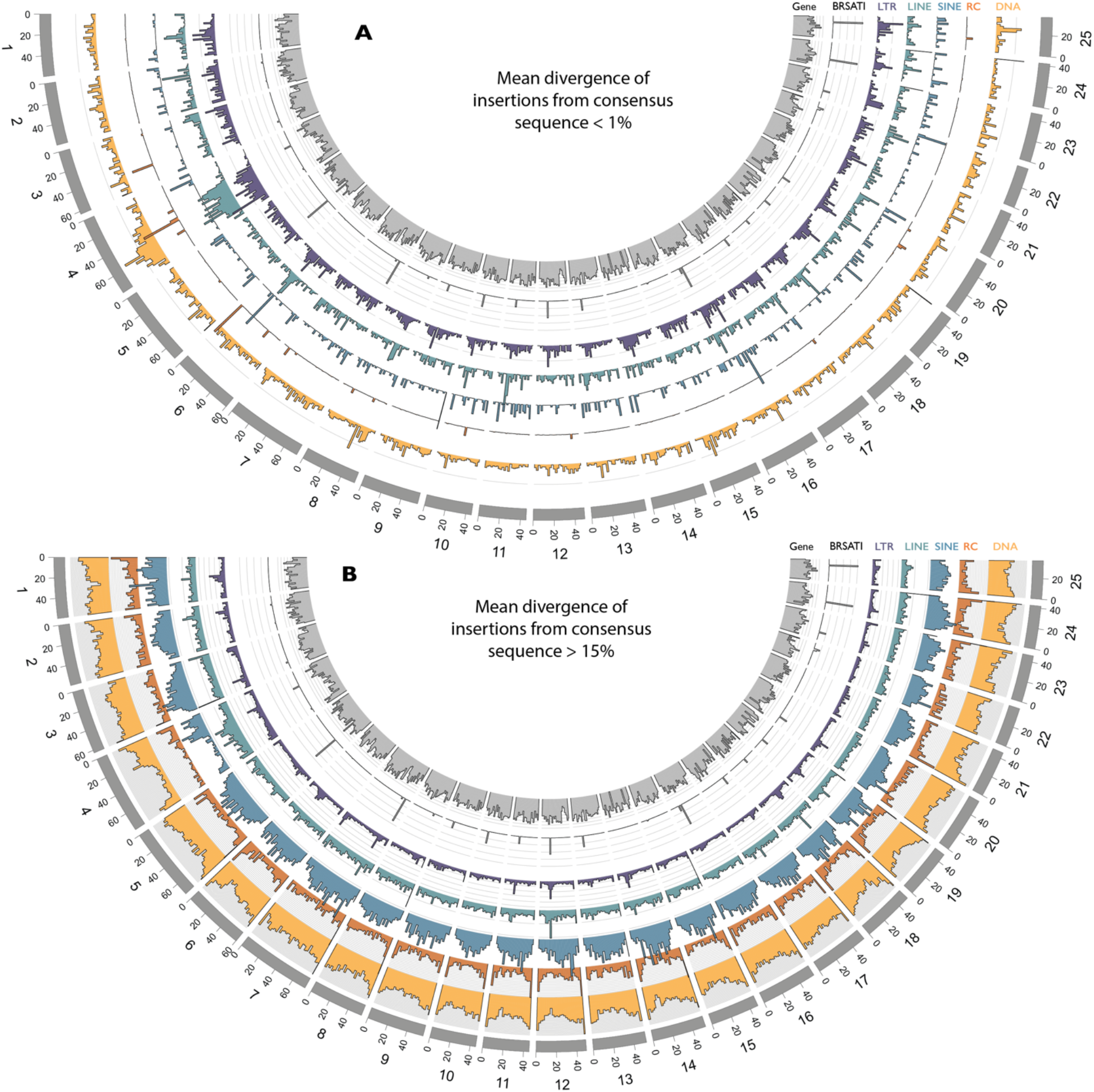
Genomic distribution of old and young insertions. Relates to figure 2A. A. Young insertions (<2% diverged from the TE family consensus sequence) are enriched in pericentromeric regions and on the long arm of chromosome 4. This is consistent with TE families preferentially targeting repetitive or heterochromatic regions. B. When restricting the analysis to older insertions (>15% diverged from consensus), the patterns are broadly similar, with the exception of chromosome 4, which is depleted, consistent with increased turnover of insertions.

**Supplementary Figure 4.**
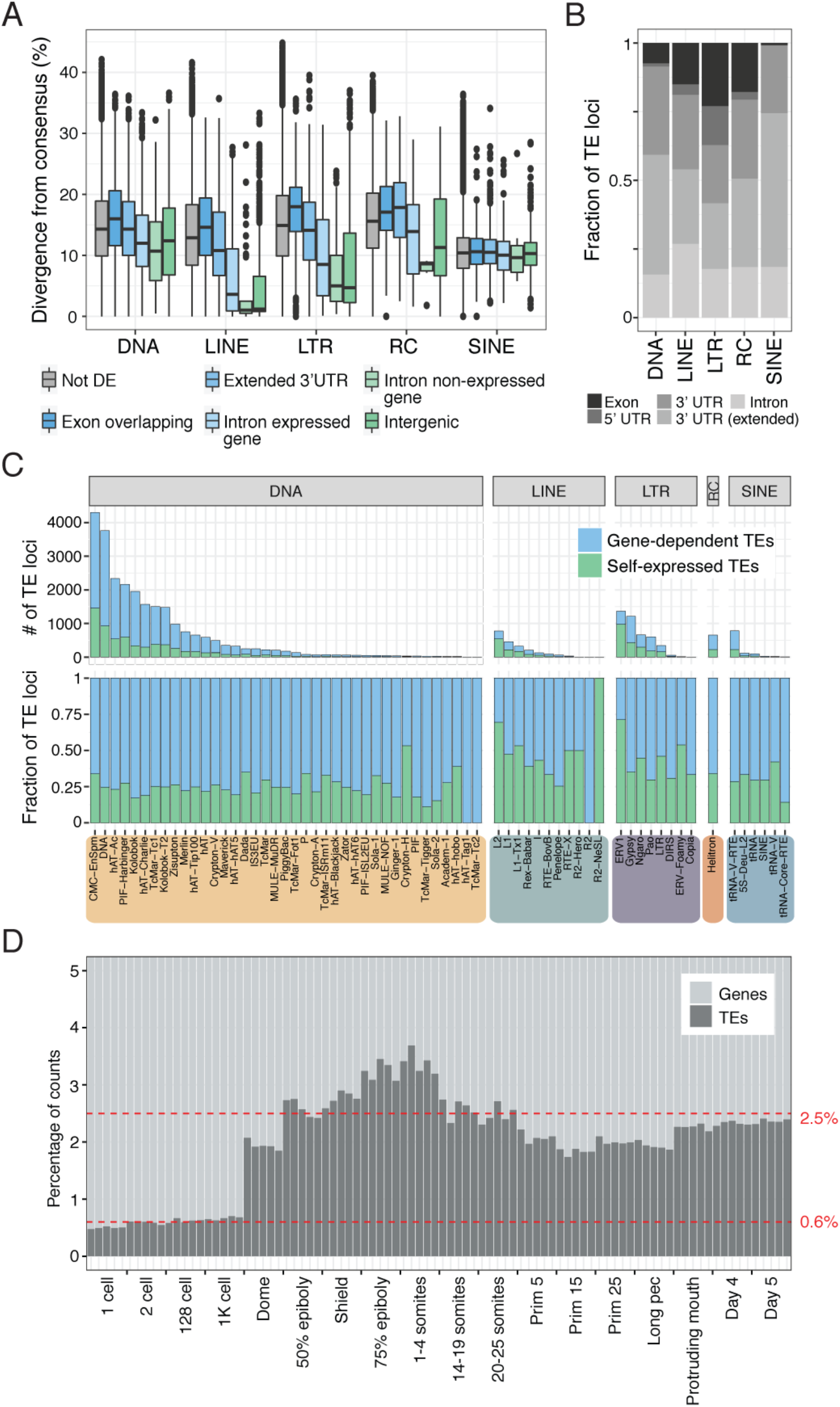
A. Boxplot of divergence from consensus sequence for differentially expressed self-expressed TEs from all TE categories. B. Fraction of differentially expressed gene-dependent TE loci contained in different gene components. C. Number (top) and fraction (bottom) of differentially expressed TE loci in gene-dependent or self-expressed categories split by TE family and TE class. D. Percentage of read counts assigned to genes or TEs for each sample. Red dashed line marks the mean of TE assigned reads for pre-ZGA (lower) and post-ZGA (upper) samples.

**Supplementary Figure 5.**
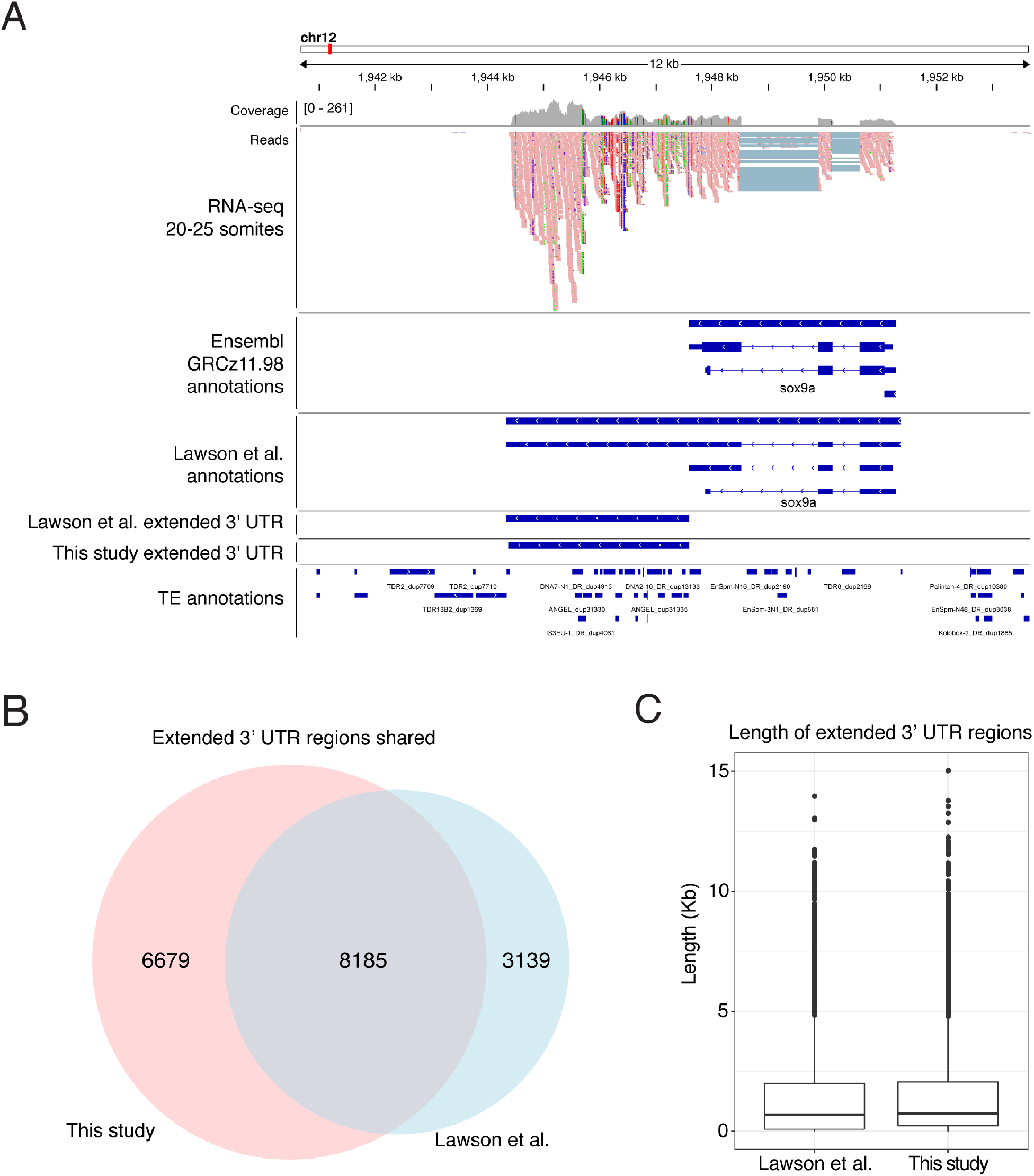
A. Genome browser snapshot of a representative locus showing un-annotated 3’ UTR region on gene sox9a. RNA-seq track shows merged replicates at 20-25 somites. Ensembl GRCz11.98 and Lawson et al. 2020 annotations are shown, together with extracted extended 3’ UTR region from Lawson et al. and this study. In this study, transcriptome assembly using StringTie was used to find extended 3’ UTR regions using White et al. 2017 RNA-seq data. B. Overlap of extended 3’ UTR regions between Lawson et al. and this study. C. Length of extended 3’ UTR regions for Lawson et al. 2020 and this study show similar distribution.

**Supplementary Figure 6.**
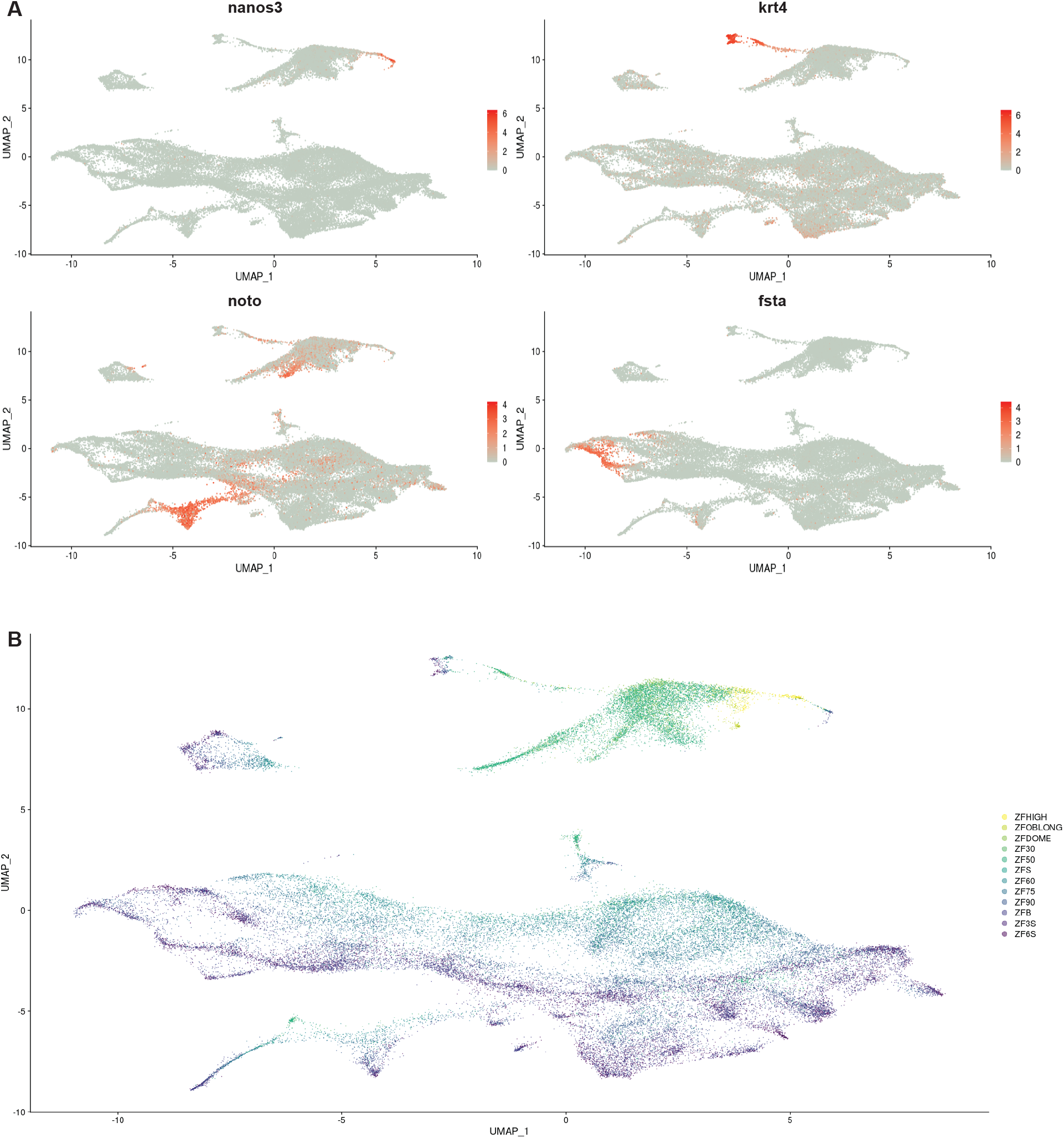
Gene markers and development stages mark the cell lineages in UMAP from single cell sequencing. A. UMAP shows the cell lineage marked by nano3, the primordial germ cell lineage; krt4, the EVL cell lineage; noto, the notochord lineage; fsta, the cephalic mesoderm lineage. B. Cells cluster together with its closet developmental stages.

**Supplementary Figure 7.**
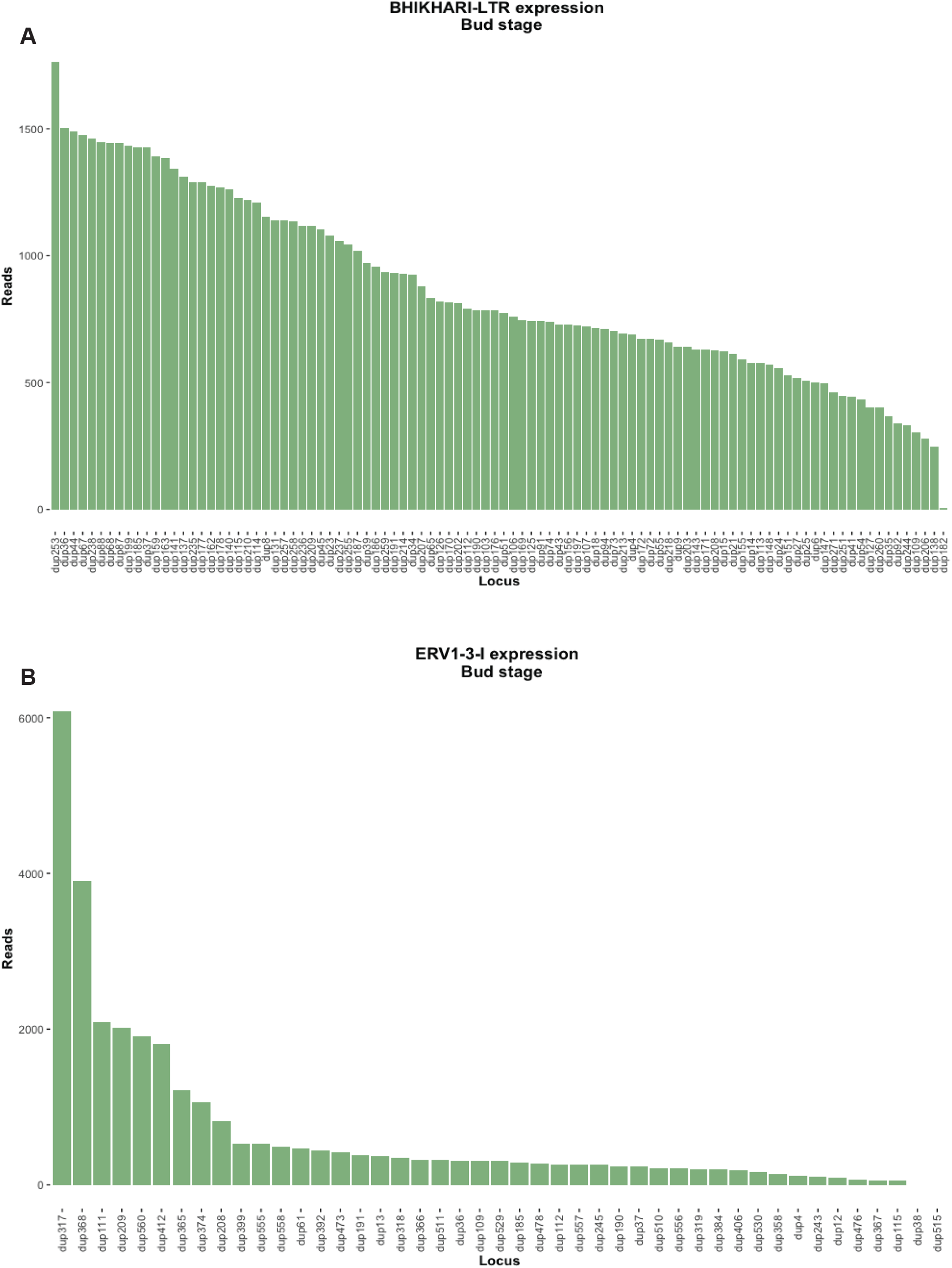
Uniquely mapping reads mapped to individual loci of A. BHIKHARI and B. ERV1-3 in Bud stage from single-cell RNA-seq. From the 98 self-expressed loci of BHIKHARI, 97 of them are expressed. Of 45 self-expressed loci of ERV1-3, 43 are expressed, but most reads are from the 10 highly expressed loci.

## References

Attig, J., Young, G.R., Hosie, L., Perkins, D., Encheva-Yokoya, V., Stoye, J.P., Snijders, A.P., Ternette, N., and Kassiotis, G. (2019). LTR retroelement expansion of the human cancer transcriptome and immunopeptidome revealed by de novo transcript assembly. Genome Res. 29, 1578–1590.

Bailly-Bechet, M., Haudry, A., and Lerat, E. (2014). “One code to find them all”: A perl tool to conveniently parse RepeatMasker output files. Mob. DNA 5, 13.

Bao, W., Kojima, K.K., and Kohany, O. (2015). Repbase Update, a database of repetitive elements in eukaryotic genomes. Mob. DNA 6, 11.

Bendall, M.L., de Mulder, M., Iñiguez, L.P., Lecanda-Sánchez, A., Pérez-Losada, M., Ostrowski, M.A., Jones, R.B., Mulder, L.C.F., Reyes-Terán, G., Crandall, K.A., et al. (2019). Telescope: Characterization of the retrotranscriptome by accurate estimation of transposable element expression. PLOS Comput. Biol. 15, e1006453.

Blanc, S., Ruggiero, F., Birot, A.-M., Acloque, H., Décimo, D., Lerat, E., Ohlmann, T., Samarut, J., and Mey, A. (2014). Subcellular Localization of ENS-1/ERNI in Chick Embryonic Stem Cells. PLoS One 9, e92039.

Blass, E., Bell, M., and Boissinot, S. (2012). Accumulation and rapid decay of non-LTR retrotransposons in the genome of the three-spine stickleback. Genome Biol. Evol. 4, 687–702.

Boissinot, S., Davis, J., Entezam, A., Petrov, D., and Furano, A. V. (2006). Fitness cost of LINE-1 (L1) activity in humans. Proc. Natl. Acad. Sci. 103, 9590–9594.

Bowen, N.J. (2003). Retrotransposons and Their Recognition of pol II Promoters: A Comprehensive Survey of the Transposable Elements From the Complete Genome Sequence of Schizosaccharomyces pombe. Genome Res. 13, 1984–1997.

Brind’Amour, J., Kobayashi, H., Richard Albert, J., Shirane, K., Sakashita, A., Kamio, A., Bogutz, A., Koike, T., Karimi, M.M., Lefebvre, L., et al. (2018). LTR retrotransposons transcribed in oocytes drive species-specific and heritable changes in DNA methylation. Nat. Commun. 9, 3331.

Bruno, M., Mahgoub, M., and Macfarlan, T.S. (2019). The Arms Race Between KRAB–Zinc Finger Proteins and Endogenous Retroelements and Its Impact on Mammals. Annu. Rev. Genet. 53, annurev-genet-112618-043717.

Chalopin, D., Naville, M., Plard, F., Galiana, D., and Volff, J.N. (2015). Comparative analysis of transposable elements highlights mobilome diversity and evolution in vertebrates. Genome Biol. Evol. 7, 567–580.

Chalvet, F., Teysset, L., Terzian, C., Prud’homme, N., Santamaria, P., Bucheton, A., and Pélisson, A. (1999). Proviral amplification of the Gypsy endogenous retrovirus of Drosophila melanogaster involves env-independent invasion of the female germline. EMBO J. 18, 2659–2669.

Chen, Y., and Schier, A.F. (2001). The zebrafish Nodal signal Squint functions as a morphogen. Nature 411, 607–610.

Chuong, E.B., Elde, N.C., and Feschotte, C. (2017). Regulatory activities of transposable elements: from conflicts to benefits. Nat. Rev. Genet. 18, 71–86.

Clayton, E.A., Rishishwar, L., Huang, T.-C., Gulati, S., Ban, D., McDonald, J.F., and Jordan, I.K. (2020). An atlas of transposable element-derived alternative splicing in cancer. Philos. Trans. R. Soc. B Biol. Sci. 375, 20190342.

Cordaux, R., Udit, S., Batzer, M.A., and Feschotte, C. (2006). Birth of a chimeric primate gene by capture of the transposase gene from a mobile element. Proc. Natl. Acad. Sci. 103, 8101–8106.

Cosby, R.L., Judd, J., Zhang, R., Zhong, A., Garry, N., Pritham, E.J., and Feschotte, C. (2021). Recurrent evolution of vertebrate transcription factors by transposase capture. Science (80-.). 371, eabc6405.

Dobin, A., Davis, C.A., Schlesinger, F., Drenkow, J., Zaleski, C., Jha, S., Batut, P., Chaisson, M., and Gingeras, T.R. (2013). STAR: ultrafast universal RNA-seq aligner. Bioinformatics 29, 15–21.

Durruthy-Durruthy, J., Sebastiano, V., Wossidlo, M., Cepeda, D., Cui, J., Grow, E.J., Davila, J., Mall, M., Wong, W.H., Wysocka, J., et al. (2016). The primate-specific noncoding RNA HPAT5 regulates pluripotency during human preimplantation development and nuclear reprogramming. Nat. Genet. 48, 44–52.

Fadloun, A., Le Gras, S., Jost, B., Ziegler-Birling, C., Takahashi, H., Gorab, E., Carninci, P., and Torres-Padilla, M.-E. (2013). Chromatin signatures and retrotransposon profiling in mouse embryos reveal regulation of LINE-1 by RNA. Nat. Struct. Mol. Biol. 20, 332–338.

Farrell, J.A., Wang, Y., Riesenfeld, S.J., Shekhar, K., Regev, A., and Schier, A.F. (2018). Single-cell reconstruction of developmental trajectories during zebrafish embryogenesis. Science (80-.). 360.

Faulkner, G.J., Kimura, Y., Daub, C.O., Wani, S., Plessy, C., Irvine, K.M., Schroder, K., Cloonan, N., Steptoe, A.L., Lassmann, T., et al. (2009). The regulated retrotransposon transcriptome of mammalian cells. Nat. Genet. 41, 563–571.

Feschotte, C., Zhang, X., and Wessler, S.R. (2002). Miniature Inverted-Repeat Transposable Elements and Their Relationship to Established DNA Transposons. In Mobile DNA II, N. Craig, R. Craigie, M. Gellert, and A. Lambowitz, eds. (Washington, DC: American Society of Microbiology), pp. 1147–1158.

Finnegan, D.J. (1989). Eukaryotic transposable elements and genome evolution. Trends Genet. 5, 103–107.

Fort, A., Hashimoto, K., Yamada, D., Salimullah, M., Keya, C.A., Saxena, A., Bonetti, A., Voineagu, I., Bertin, N., Kratz, A., et al. (2014). Deep transcriptome profiling of mammalian stem cells supports a regulatory role for retrotransposons in pluripotency maintenance. Nat. Genet. 46, 558–566.

Frahry, M.B., Sun, C., Chong, R.A., and Mueller, R.L. (2015). Low Levels of LTR Retrotransposon Deletion by Ectopic Recombination in the Gigantic Genomes of Salamanders. J. Mol. Evol. 80, 120–129.

Franke, V., Ganesh, S., Karlic, R., Malik, R., Pasulka, J., Horvat, F., Kuzman, M., Fulka, H., Cernohorska, M., Urbanova, J., et al. (2017). Long terminal repeats power evolution of genes and gene expression programs in mammalian oocytes and zygotes. Genome Res. 27, 1384–1394.

Fricker, A.D., and Peters, J.E. (2014). Vulnerabilities on the Lagging-Strand Template: Opportunities for Mobile Elements. Annu. Rev. Genet. 48, 167–186.

Furano, A. V., Duvernell, D.D., and Boissinot, S. (2004). L1 (LINE-1) retrotransposon diversity differs dramatically between mammals and fish. Trends Genet. 20, 9–14.

Gao, B., Shen, D., Xue, S., Chen, C., Cui, H., and Song, C. (2016). The contribution of transposable elements to size variations between four teleost genomes. Mob. DNA 7, 4.

Garcia-Perez, J.L., Widmann, T.J., and Adams, I.R. (2016). The impact of transposable elements on mammalian development. Development 143, 4101–4114.

Gibbs, R.A., Weinstock, G.M., Metzker, M.L., Muzny, D.M., Sodergren, E.J., Scherer, S., Scott, G., Steffen, D., Worley, K.C., Burch, P.E., et al. (2004). Genome sequence of the Brown Norway rat yields insights into mammalian evolution. Nature 428, 493–521.

Göke, J., Lu, X., Chan, Y.-S., Ng, H.-H., Ly, L.-H., Sachs, F., and Szczerbinska, I. (2015). Dynamic Transcription of Distinct Classes of Endogenous Retroviral Elements Marks Specific Populations of Early Human Embryonic Cells. Cell Stem Cell 16, 135–141.

Grow, E.J., Flynn, R.A., Chavez, S.L., Bayless, N.L., Wossidlo, M., Wesche, D.J., Martin, L., Ware, C.B., Blish, C.A., Chang, H.Y., et al. (2015). Intrinsic retroviral reactivation in human preimplantation embryos and pluripotent cells. Nature 522, 221–225.

Haig, D. (2016). Transposable elements: Self-seekers of the germline, team-players of the soma. BioEssays 38, 1158–1166.

He, J., Babarinde, I.A., Sun, L., Xu, S., Chen, R., Shi, J., Wei, Y., Li, Y., Ma, G., Zhuang, Q., et al. (2021). Identifying transposable element expression dynamics and heterogeneity during development at the single-cell level with a processing pipeline scTE. Nat. Commun. 12, 1456.

Hermant, C., and Torres-Padilla, M.-E. (2021). TFs for TEs: the transcription factor repertoire of mammalian transposable elements.

Heyn, P., Kircher, M., Dahl, A., Kelso, J., Tomancak, P., Kalinka, A.T., and Neugebauer, K.M. (2014). The Earliest Transcribed Zygotic Genes Are Short, Newly Evolved, and Different across Species. Cell Rep. 6, 285–292.

Hickman, A.B., and Dyda, F. (2016). DNA Transposition at Work. Chem. Rev. 116, 12758–12784.

Houwing, S., Kamminga, L.M., Berezikov, E., Cronembold, D., Girard, A., van den Elst, H., Filippov, D. V, Blaser, H., Raz, E., Moens, C.B., et al. (2007). A Role for Piwi and piRNAs in Germ Cell Maintenance and Transposon Silencing in Zebrafish. Cell 129, 69–82.

Houwing, S., Berezikov, E., and Ketting, R.F. (2008). Zili is required for germ cell differentiation and meiosis in zebrafish. EMBO J. 27, 2702–2711.

Howe, K. (2020). The zebrafish genome sequencing project: bioinformatics resources. In Behavioral and Neural Genetics of Zebrafish, (Elsevier), pp. 551–562.

Howe, K., Clark, M.D., Torroja, C.F., Torrance, J., Berthelot, C., Muffato, M., Collins, J.J.E., Humphray, S., McLaren, K., Matthews, L., et al. (2013). The zebrafish reference genome sequence and its relationship to the human genome. Nature 496, 498–503.

Ishiuchi, T., Enriquez-Gasca, R., Mizutani, E., Bošković, A., Ziegler-Birling, C., Rodriguez-Terrones, D., Wakayama, T., Vaquerizas, J.M., and Torres-Padilla, M.-E. (2015). Early embryonic-like cells are induced by downregulating replication-dependent chromatin assembly. Nat. Struct. Mol. Biol. 22, 662–671.

Jachowicz, J.W., Bing, X., Pontabry, J., Bošković, A., Rando, O.J., and Torres-Padilla, M.-E. (2017). LINE-1 activation after fertilization regulates global chromatin accessibility in the early mouse embryo. Nat. Genet. 49, 1502–1510.

Jin, Y., Tam, O.H., Paniagua, E., and Hammell, M. (2015). TEtranscripts: A package for including transposable elements in differential expression analysis of RNA-seq datasets. Bioinformatics 31, 3593–3599.

Kaaij, L.J.T., Hoogstrate, S.W., Berezikov, E., and Ketting, R.F. (2013). piRNA dynamics in divergent zebrafish strains reveal long-lasting maternal influence on zygotic piRNA profiles. RNA 19, 345–356.

Kapusta, A., Kronenberg, Z., Lynch, V.J., Zhuo, X., Ramsay, L., Bourque, G., Yandell, M., and Feschotte, C. (2013). Transposable Elements Are Major Contributors to the Origin, Diversification, and Regulation of Vertebrate Long Noncoding RNAs. PLoS Genet. 9, e1003470.

Kapusta, A., Suh, A., and Feschotte, C. (2017). Dynamics of genome size evolution in birds and mammals. Proc. Natl. Acad. Sci. 114, E1460–E1469.

Katoh, K., and Standley, D.M. (2013). MAFFT Multiple Sequence Alignment Software Version 7: Improvements in Performance and Usability. Mol. Biol. Evol. 30, 772–780.

Kelley, D., and Rinn, J. (2012). Transposable elements reveal a stem cell-specific class of long noncoding RNAs. Genome Biol. 13, R107.

Kong, Y., Rose, C.M., Cass, A.A., Williams, A.G., Darwish, M., Lianoglou, S., Haverty, P.M., Tong, A.-J., Blanchette, C., Albert, M.L., et al. (2019). Transposable element expression in tumors is associated with immune infiltration and increased antigenicity. Nat. Commun. 10, 5228.

Krzywinski, M., Schein, J., Birol, I., Connors, J., Gascoyne, R., Horsman, D., Jones, S.J., and Marra, M.A. (2009). Circos: An information aesthetic for comparative genomics. Genome Res. 19, 1639–1645.

Kung, J.T.Y., Colognori, D., and Lee, J.T. (2013). Long Noncoding RNAs: Past, Present, and Future. Genetics 193, 651–669.

Lanciano, S., and Cristofari, G. (2020). Measuring and interpreting transposable element expression. Nat. Rev. Genet.

Lander, E.S., Linton, L.M., Birren, B., Nusbaum, C., Zody, M.C., Baldwin, J., Devon, K., Dewar, K., Doyle, M., FitzHugh, W., et al. (2001). Initial sequencing and analysis of the human genome. Nature 409, 860–921.

Langmead, B., and Salzberg, S.L. (2012). Fast gapped-read alignment with Bowtie 2. Nat. Methods 9, 357–359.

Lawson, N.D., Li, R., Shin, M., Grosse, A., Yukselen, O., Stone, O.A., Kucukural, A., and Zhu, L. (2020). An improved zebrafish transcriptome annotation for sensitive and comprehensive detection of cell type-specific genes. Elife 9.

Lev-Maor, G., Sorek, R., Shomron, N., and Ast, G. (2003). The birth of an alternatively spliced exon: 3’ splice-site selection in Alu exons. Science 300, 1288–1291.

Li, H., Handsaker, B., Wysoker, A., Fennell, T., Ruan, J., Homer, N., Marth, G., Abecasis, G., Durbin, R., and 1000 Genome Project Data Processing Subgroup (2009). The Sequence Alignment/Map format and SAMtools. Bioinformatics 25, 2078–2079.

Liu, J., Zhu, Y., Luo, G.Z., Wang, X., Yue, Y., Wang, X., Zong, X., Chen, K., Yin, H., Fu, Y., et al. (2016). Abundant DNA 6mA methylation during early embryogenesis of zebrafish and pig. Nat. Commun. 7, 1–7.

Love, M.I., Huber, W., and Anders, S. (2014). Moderated estimation of fold change and dispersion for RNA-seq data with DESeq2. Genome Biol. 15, 550.

Lu, X., Sachs, F., Ramsay, L., Jacques, P.-É., Göke, J., Bourque, G., and Ng, H.-H. (2014). The retrovirus HERVH is a long noncoding RNA required for human embryonic stem cell identity. Nat. Struct. Mol. Biol. 21, 423–425.

Luo, R., An, M., Arduini, B.L., and Henion, P.D. (2001). Specific pan-neural crest expression of zebrafish Crestin throughout embryonic development. Dev. Dyn. 220, 169–174.

Macfarlan, T.S., Gifford, W.D., Driscoll, S., Lettieri, K., Rowe, H.M., Bonanomi, D., Firth, A., Singer, O., Trono, D., and Pfaff, S.L. (2012). Embryonic stem cell potency fluctuates with endogenous retrovirus activity. Nature 487, 57–63.

Macosko, E.Z., Basu, A., Satija, R., Nemesh, J., Shekhar, K., Goldman, M., Tirosh, I., Bialas, A.R., Kamitaki, N., Martersteck, E.M., et al. (2015). Highly Parallel Genome-wide Expression Profiling of Individual Cells Using Nanoliter Droplets. Cell 161, 1202–1214.

Medstrand, P., Van De Lagemaat, L.N., and Mager, D.L. (2002). Retroelement distributions in the human genome: Variations associated with age and proximity to genes. Genome Res. 12, 1483–1495.

Newman, J.C., Bailey, A.D., Fan, H.-Y., Pavelitz, T., and Weiner, A.M. (2008). An Abundant Evolutionarily Conserved CSB-PiggyBac Fusion Protein Expressed in Cockayne Syndrome. PLoS Genet. 4, e1000031.

Ng, K.W., Attig, J., Bolland, W., Young, G.R., Major, J., Wrobel, A.G., Gamblin, S., Wack, A., and Kassiotis, G. (2020). Tissue-specific and interferon-inducible expression of nonfunctional ACE2 through endogenous retroelement co-option. Nat. Genet. 52, 1294–1302.

Pace, J.K., and Feschotte, C. (2007). The evolutionary history of human DNA transposons: Evidence for intense activity in the primate lineage. Genome Res. 17, 422–432.

Palazzo, A., Caizzi, R., Viggiano, L., and Marsano, R.M. (2017). Does the Promoter Constitute a Barrier in the Horizontal Transposon Transfer Process? Insight from Bari Transposons. Genome Biol. Evol. 9, 1637–1645.

Palazzo, A., Lorusso, P., Miskey, C., Walisko, O., Gerbino, A., Marobbio, C.M.T., Ivics, Z., and Marsano, R.M. (2019). Transcriptionally promiscuous “blurry” promoters in Tc1/mariner transposons allow transcription in distantly related genomes. Mob. DNA 10, 13.

Peaston, A.E., Evsikov, A. V., Graber, J.H., de Vries, W.N., Holbrook, A.E., Solter, D., and Knowles, B.B. (2004). Retrotransposons Regulate Host Genes in Mouse Oocytes and Preimplantation Embryos. Dev. Cell 7, 597–606.

Percharde, M., Lin, C.-J., Yin, Y., Guan, J., Peixoto, G.A., Bulut-Karslioglu, A., Biechele, S., Huang, B., Shen, X., and Ramalho-Santos, M. (2018). A LINE1-Nucleolin Partnership Regulates Early Development and ESC Identity. Cell 174, 391–405.e19.

Pertea, M., Pertea, G.M., Antonescu, C.M., Chang, T.-C., Mendell, J.T., and Salzberg, S.L. (2015). StringTie enables improved reconstruction of a transcriptome from RNA-seq reads. Nat. Biotechnol. 33, 290–295.

Petri, R., Brattås, P.L., Sharma, Y., Jönsson, M.E., Pircs, K., Bengzon, J., and Jakobsson, J. (2019). LINE-2 transposable elements are a source of functional human microRNAs and target sites. PLoS Genet. 15, e1008036.

Petrov, D.A., Aminetzach, Y.T., Davis, J.C., Bensasson, D., and Hirsh, A.E. (2003). Size matters: Non-LTR retrotransposable elements and ectopic recombination in drosophila. Mol. Biol. Evol. 20, 880–892.

Petrov, D.A., Fiston-Lavier, A.S., Lipatov, M., Lenkov, K., and González, J. (2011). Population genomics of transposable elements in drosophila melanogaster. Mol. Biol. Evol. 28, 1633–1644.

Phillips, R.B., and Reed, K.M. (2000). Localization of repetitive DNAs to zebrafish (Danio rerio) chromosomes by fluorescence in situ hybridization (FISH). Chromosom. Res. 8, 27–35.

Platt, R.N., Mangum, S.F., and Ray, D.A. (2016). Pinpointing the vesper bat transposon revolution using the Miniopterus natalensis genome. Mob. DNA 7, 12.

Price, M.N., Dehal, P.S., and Arkin, A.P. (2010). FastTree 2 – Approximately Maximum-Likelihood Trees for Large Alignments. PLoS One 5, e9490.

Pritham, E.J., and Feschotte, C. (2007). Massive amplification of rolling-circle transposons in the lineage of the bat Myotis lucifugus. Proc. Natl. Acad. Sci. 104, 1895–1900.

Quinlan, A.R., and Hall, I.M. (2010). BEDTools: a flexible suite of utilities for comparing genomic features. Bioinformatics 26, 841–842.

Raz, E. (2003). Primordial germ-cell development: the zebrafish perspective. Nat. Rev. Genet. 4, 690–700.

Richardson, S.R., Gerdes, P., Gerhardt, D.J., Sanchez-Luque, F.J., Bodea, G.-O., Muñoz-Lopez, M., Jesuadian, J.S., Kempen, M.-J.H.C., Carreira, P.E., Jeddeloh, J.A., et al. (2017). Heritable L1 retrotransposition in the mouse primordial germline and early embryo. Genome Res. 27, 1395–1405.

Risso, D., Ngai, J., Speed, T.P., and Dudoit, S. (2014). Normalization of RNA-seq data using factor analysis of control genes or samples. Nat. Biotechnol. 32, 896–902.

Robbez-Masson, L., and Rowe, H.M. (2015). Retrotransposons shape species-specific embryonic stem cell gene expression. Retrovirology 12, 45.

Rodriguez-Terrones, D., and Torres-Padilla, M.-E. (2018). Nimble and Ready to Mingle: Transposon Outbursts of Early Development Box 1. Transposable Elements Exhibit Limited Conservation across Mammals. Trends Genet. 34.

Romanish, M.T., Lock, W.M., de Lagemaat, L.N. van, Dunn, C.A., and Mager, D.L. (2007). Repeated Recruitment of LTR Retrotransposons as Promoters by the Anti-Apoptotic Locus NAIP during Mammalian Evolution. PLoS Genet. 3, e10.

Rubinstein, A.L., Lee, D., Luo, R., Henion, P.D., and Halpern, M.E. (2000). Genes dependent on zebrafish cyclops function identified by AFLP differential gene expression screen. Genesis 26, 86–97.

Shao, W., and Wang, T. (2021). Transcript assembly improves expression quantification of transposable elements in single-cell RNA-seq data. Genome Res. 31, 88–100.

Smit, A.F.A., Hubley, R., and Green, P. RepeatMasker Open-4.0.

Sotero-Caio, C.G., Platt, R.N., Suh, A., and Ray, D.A. (2017). Evolution and diversity of transposable elements in vertebrate genomes. Genome Biol. Evol. 9, 161–177.

Spradling, A.C., Bellen, H.J., and Hoskins, R.A. (2011). Drosophila P elements preferentially transpose to replication origins. Proc. Natl. Acad. Sci. 108, 15948–15953.

Stitzer, M.C., Anderson, S.N., Springer, N.M., and Ross-Ibarra, J. (2019). The Genomic Ecosystem of Transposable Elements in Maize. BioRxiv.

Storer, J., Hubley, R., Rosen, J., Wheeler, T.J., and Smit, A.F. (2021). The Dfam community resource of transposable element families, sequence models, and genome annotations. Mob. DNA 12, 2.

Svoboda, P., Stein, P., Anger, M., Bernstein, E., Hannon, G.J., and Schultz, R.M. (2004). RNAi and expression of retrotransposons MuERV-L and IAP in preimplantation mouse embryos. Dev. Biol. 269, 276–285.

Tarasov, A., Vilella, A.J., Cuppen, E., Nijman, I.J., and Prins, P. (2015). Sambamba: fast processing of NGS alignment formats. Bioinformatics 31, 2032–2034.

Thisse, C., and Thisse, B. (2008). High-resolution in situ hybridization to whole-mount zebrafish embryos. Nat. Protoc. 3, 59–69.

Thomas, J., Schaack, S., and Pritham, E.J. (2010). Pervasive Horizontal Transfer of Rolling-Circle Transposons among Animals. Genome Biol. Evol. 2, 656–664.

Thompson, P.J., Macfarlan, T.S., and Lorincz, M.C. (2016). Long Terminal Repeats: From Parasitic Elements to Building Blocks of the Transcriptional Regulatory Repertoire. Mol. Cell 62, 766–776.

Topczewska, J.M. (2001). The winged helix transcription factor Foxc1a is essential for somitogenesis in zebrafish. Genes Dev. 15, 2483–2493.

Vogel, A.M., and Gerster, T. (1999). Promoter activity of the zebrafish bhikhari retroelement requires an intact activin signaling pathway. Mech. Dev. 85, 133–146.

Wang, J., Xie, G., Singh, M., Ghanbarian, A.T., Raskó, T., Szvetnik, A., Cai, H., Besser, D., Prigione, A., Fuchs, N. V., et al. (2014). Primate-specific endogenous retrovirus-driven transcription defines naive-like stem cells. Nature 516, 405–409.

Wang, J., Singh, M., Sun, C., Besser, D., Prigione, A., Ivics, Z., Hurst, L.D., and Izsvák, Z. (2016). Isolation and cultivation of naive-like human pluripotent stem cells based on HERVH expression. Nat. Protoc. 11, 327–346.

Wang, L., Dou, K., Moon, S., Tan, F.J., and Zhang, Z.Z. (2018). Hijacking Oogenesis Enables Massive Propagation of LINE and Retroviral Transposons. Cell 174, 1082–1094.e12.

Waterston, R.H., Lindblad-Toh, K., Birney, E., Rogers, J., Abril, J.F., Agarwal, P., Agarwala, R., Ainscough, R., Alexandersson, M., An, P., et al. (2002). Initial sequencing and comparative analysis of the mouse genome. Nature 420, 520–562.

Wells, J.N., and Feschotte, C. (2020). A Field Guide to Eukaryotic Transposable Elements. Annu. Rev. Genet. 54, 539–561.

White, R.J., Collins, J.E., Sealy, I.M., Wali, N., Dooley, C.M., Digby, Z., Stemple, D.L., Murphy, D.N., Billis, K., Hourlier, T., et al. (2017). A high-resolution mRNA expression time course of embryonic development in zebrafish. Elife 6.

Wicker, T., Sabot, F., Hua-Van, A., Bennetzen, J.L., Capy, P., Chalhoub, B., Flavell, A., Leroy, P., Morgante, M., Panaud, O., et al. (2007). A unified classification system for eukaryotic transposable elements. Nat. Rev. Genet. 8, 973–982.

Wilm, B., James, R.G., Schultheiss, T.M., and Hogan, B.L.M. (2004). The forkhead genes, Foxc1 and Foxc2, regulate paraxial versus intermediate mesoderm cell fate. Dev. Biol. 271, 176–189.

Yang, H., Luan, Y., Liu, T., Lee, H.J., Fang, L., Wang, Y., Wang, X., Zhang, B., Jin, Q., Ang, K.C., et al. (2020). A map of cis-regulatory elements and 3D genome structures in zebrafish. Nature.

Yu, G., Wang, L.-G., and He, Q.-Y. (2015). ChIPseeker: an R/Bioconductor package for ChIP peak annotation, comparison and visualization. Bioinformatics 31, 2382–2383.

Zaghlool, A., Ameur, A., Nyberg, L., Halvardson, J., Grabherr, M., Cavelier, L., and Feuk, L. (2013). Efficient cellular fractionation improves RNA sequencing analysis of mature and nascent transcripts from human tissues. BMC Biotechnol. 13, 99.

Zarnack, K., König, J., Tajnik, M., Martincorena, I., Eustermann, S., Stévant, I., Reyes, A., Anders, S., Luscombe, N.M., and Ule, J. (2013). Direct competition between hnRNP C and U2AF65 protects the transcriptome from the exonization of Alu elements. Cell 152, 453–466.

