## Supplementary Data 1 for "A genomic portrait of zebrafish transposable elements and their spatiotemporal embryonic expression": README

# Supplementary Data 1

This folder contains the raw repeatmasker output file, danRer11.nonalt.fa.out,
along with a summary file, danRer11.nonalt.fa.tbl. The parserm folder contains
the output from Aurelie Kapusta's ParseRM.pl script
(https://github.com/4ureliek/Parsing-RepeatMasker-Outputs/blob/master/parseRM.pl),
which provides more accurate figures of TE composition and age than the default
repeatmasker output.
