## Supplementary Data 3 for "A genomic portrait of zebrafish transposable elements and their spatiotemporal embryonic expression": README.txt

Column header descriptions

tename: family name of TE as given in Repbase/DFAM
flatname: family name with underscores converted to hyphens
mergedname: LTR and internal regions given single name
tefam: superfamily of TE
teclass: class of TE
cov: percent genome coverage (calculated from parseRM.pl)
frags: number of masked fragments (from parseRM.pl)
copynum: TE copynumber, estimated from defragmented insertions
length: length of TE consensus sequence (combining LTRs and internal)
meandiv: meand divergence of insertions from consensus sequence
medlen: median terminal branch length of insertions on phylo tree
meddist: median distance from nearest genes
ssdist: median distance from genes on same strand
dsdist: mediatn distance from genes on opposite strand
meantau: mean scRNA cell-type specificity tau value (mean across stages)
sense/anti: scRNA sense/antisense ratio
robust: True if scRNA expressed in at least 5% of cells in at least 1 cluster
isltr: True if LTR region (as opposed to internal)
